# Structure of a monomeric photosystem I core associated with iron-stress-induced-A proteins from *Anabaena* sp. PCC 7120

**DOI:** 10.1101/2022.08.05.501323

**Authors:** Ryo Nagao, Koji Kato, Tasuku Hamaguchi, Yoshifumi Ueno, Naoki Tsuboshita, Shota Shimizu, Miyu Furutani, Shigeki Ehira, Yoshiki Nakajima, Keisuke Kawakami, Takehiro Suzuki, Naoshi Dohmae, Seiji Akimoto, Koji Yonekura, Jian-Ren Shen

## Abstract

Iron-stress-induced-A proteins (IsiAs) are expressed in cyanobacteria under iron-deficient conditions. The cyanobacterium *Anabaena* sp. PCC 7120 has four *isiA* genes; however, their binding property and functional roles in PSI are still missing. We analyzed a cryo-electron microscopy structure of a PSI-IsiA supercomplex isolated from *Anabaena* grown under an iron-deficient condition. The PSI-IsiA structure contains six IsiA subunits associated with the PsaA side of a PSI core monomer. Three of the six IsiA subunits are identified as IsiA1 and IsiA2. The PSI-IsiA structure lacks a PsaL subunit; instead, a C-terminal domain of IsiA2 is inserted at the position of PsaL, which inhibits the oligomerization of PSI, leading to the formation of a monomer. Furthermore, excitation-energy transfer from IsiAs to PSI appeared with a time constant of 55 ps. These findings provide novel insights into both the molecular assembly of the *Anabaena* IsiA family and the functional roles of IsiAs.

---

Oxygenic photosynthesis of cyanobacteria, various algae, and land plants converts light energy from the sun into biologically useful chemical energy concomitant with the evolution of molecular oxygen^1^. The central part of the light-energy conversion is two multi-subunit pigment-protein complexes, photosystem I and photosystem II (PSI and PSII, respectively), which perform light-driven charge separation and a series of electron transfer reactions^1^. Among these complexes, PSII organizes mainly into a dimer regardless of species of the organism^2,3^, whereas PSI exhibits different structural organization among photosynthetic organisms^4–6^. Prokaryotic cyanobacteria have trimeric^7–10^ or tetrameric PSI^10–14^ in addition to other minor forms of PSI monomers and dimers^15,16^.

Light-harvesting complexes (LHCs) are responsible for harvesting light energy which is transferred to the two photosystem cores to initiate the charge separation and electron transfer reactions^1^. LHCs are mainly separated into two groups: membrane-embedded LHCs and water-soluble LHCs^17^. Iron-stress-induced-A proteins (IsiAs) are one of the membrane-embedded LHCs^18–20^ and are expressed under iron-starvation conditions in various cyanobacteria^19,20^. Structural studies have revealed that up to 18 copies of the IsiA subunit surround a trimeric PSI core, forming a PSI-IsiA supercomplex with a closed ring of IsiAs^21–25^, where IsiA can function to donate excitation energy to PSI^25–28^.

A very attractive feature of the IsiA family is that the number of *isiA* genes differs among cyanobacteria. The cyanobacterium *Leptolyngbya* sp. strain JSC-1 (hereafter referred to as *Leptolyngbya)* has five *isiA* genes *isiA1, isiA2, isiA3, isiA4,* and *isiA5,* all of which were expressed under an iron-deficient condition^29^. Our phylogenetic analysis proposed that the cyanobacterium *Anabaena* sp. PCC 7120 (hereafter referred to as *Anabaena)* has four types of *isiA* genes: *isiA1, isiA2, isiA3,* and *isiA5,* based on the similarities of sequences to *Leptolyngbya*^30^. Our previous study also showed that the IsiA forming the ring structures surrounding the PSI core trimer in *Synechocystis* sp. PCC 6803, *Synechococcus elongatus* PCC 7942, and *Thermosynechococcus vulcanus* NIES-2134^23–25^ belongs to *isiA1* of *Anabaena*^30^. Immunological analysis of IsiAs detected one band in thylakoids of *Anabaena* prepared from the iron-deficient cells^30^. In addition, two-dimensional blue-native (BN)/SDS-PAGE analysis together with mass spectrometry detected the IsiA1 subunit in a PSI fraction located near the band of the PSII dimer, showing the formation of a PSI-IsiA1 supercomplex in *Anabaena*^30^. Since *Anabaena* has not only tetrameric PSI cores^11–13^ but also PSI monomers and dimers^15^, it was suggested that the supercomplex was composed of either a PSI core monomer with several IsiA1 subunits or a PSI dimer with a few IsiA1s based on the putative molecular weight of PSI-IsiA1^30^. On the other hand, the corresponding bands of IsiA2, IsiA3, and IsiA5 subunits could not be found on the gel of the PSI-IsiA1 fraction^30^. These observations raise three issues as to (1) how the PSI-IsiA supercomplex is organized, (2) why the IsiA subunits are not associated with the PSI tetramer, and (3) whether the *isiA2, isiA3,* and *isiA5* genes are expressed or not.

To clarify these three issues, we solved a structure of a PSI-IsiA supercomplex purified from *Anabaena* grown under the iron-deficient condition by cryo-electron microscopy (cryo-EM) single-particle analysis. The results showed that PSI-IsiA supercomplex consists of six IsiA subunits encoded by different *isiA* genes associated at one site of a monomeric PSI core, forming an unclosed, monomeric PSI-IsiA supercomplex. The expression of the different *isiA* genes, the association pattern of each IsiA with the PSI core, together with their roles in energy transfer are revealed and discussed.

## Results

### Expression and accumulation of IsiAs

The expression of the *isiA* genes in *Anabaena* was examined by qRT-PCR (Supplementary Fig. 1a), which showed that the transcript levels of three *isiA* genes, *isiA1, isiA3,* and *isiA5,* are markedly increased under the iron-deficient condition, whereas the transcript level of *isiA2* is increased to a less remarkable level. The PSI-IsiA supercomplexes were purified by trehalose density gradient centrifugation (Supplementary Fig. 1b; see Methods), which showed that the supercomplex contains all of the four IsiAs as detected by mass spectrometry analysis (Supplementary Fig. 1c). The molecular weight of the purified PSI-IsiA is close to the PSI dimer purified from the strain grown under an iron-replete condition (Supplementary Fig. 1d). This oligomeric state of PSI-IsiA was also observed by BN-PAGE in thylakoid membranes isolated from *Anabaena* grown under the iron-deficient condition^31^. UV-vis spectroscopic analyses and pigment compositions of the purified supercomplex are summarized in Supplementary Fig. 1e–g, which showed that the supercomplex is characteristic of a PSI-like preparation in terms of the absorption and fluorescence bands, and contains chlorophylls (Chls), *β-* carotenes and a small amount of echinenones.

### Overall structure of the PSI-IsiA supercomplex

Cryo-EM images of the PSI-IsiA supercomplex were obtained by a JEOL CRYO ARM 300 electron microscope operated at 300 kV. After processing of the images with RELION (Supplementary Fig. 2, Supplementary Table 1), the final cryo-EM map was determined with a C1 symmetry at a resolution of 2.62 Å, based on the “gold standard” Fourier shell correlation (FSC) = 0.143 criterion (Supplementary Fig. 3).

The atomic model of PSI-IsiA was built based on the cryo-EM maps (see Methods; Fig. 1, Supplementary Table 1–3), which revealed a PSI monomeric core associated with six unique subunits outside of PsaA (Fig. 1a, b). Five of the six subunits have six membrane-spanning helices that bind Chls and carotenoids (Cars), which are characteristic of IsiAs^23–25^. The remaining subunit has nine membrane-spanning helices, six of which are similar to the former five subunits. Thus, they are assigned to IsiAs, which are numbered from 1 to 6 (Fig. 1a). Only three subunits of IsiAs are identified as IsiA1 at positions 4/5 and as IsiA2 at position 1, which are named as IsiA1-4, IsiA1-5, and IsiA2-1. The remaining three subunits IsiA-2, IsiA-3, and IsiA-6 were not identified and therefore modeled as polyalanines based on their positions (Fig. 1a). The cofactors in the IsiA subunits are summarized in Supplementary Table 3.

**Fig. 1.**
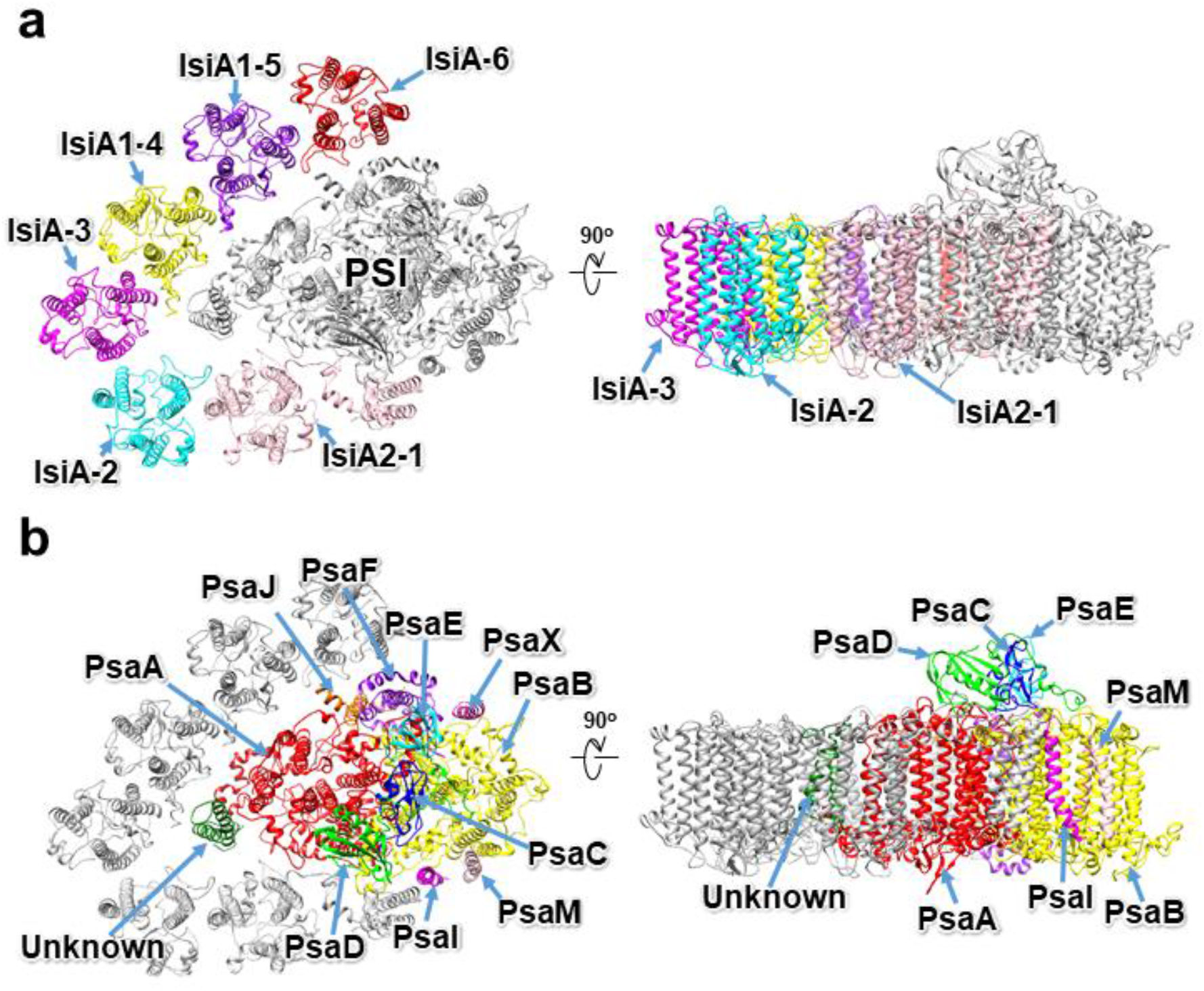
Overall structure of the PSI-IsiA supercomplex from *Anabaena*. Structures are viewed from the cytosolic side (left panels) and the direction perpendicular to the membrane normal (right panels). Only protein structures are shown, whereas those of cofactors are omitted for clarity. The IsiA (**a**) and PSI core (**b**) subunits are labeled with different colors.

### Structure of the PSI monomer

The overall architecture of the PSI core in the PSI-IsiA supercomplex is similar to that of the PSI tetramer isolated from *Anabaena*^11–13^. The PSI monomer contains 11 subunits, ten of which are PsaA, PsaB, PsaC, PsaD, PsaE, PsaF, PsaI, PsaJ, PsaM, and PsaX (Fig. 1b). The remaining one subunit is located at the position corresponding to PsaK, which was modeled as polyalanines (Supplementary Fig. 4a). This cyanobacterium has the *psaK* gene in addition to two unique genes of *alr5290* and *asr5289* with sequences similar to *psaK.* The amino acid sequences of Alr5290 and Asr5289 have similarities of 40% and 38% with that of PsaK (Supplementary Fig. 4b). However, none of the three sequences can be fitted into the density of the map. This subunit is therefore named Unknown in the present structure. Since PsaK has been solved in the *Anabaena* PSI structure under iron-replete conditions^11–13^, it seems that the cells grown under iron-deficient conditions induce the substitution of PsaK with Unknown or inhibit the expression of PsaK. Very interestingly, PsaL is lacking in the PSI-IsiA structure; instead, a part of IsiA2-1 is located at the position of PsaL (Supplementary Fig. 5a). The superposition of the PSI-IsiA structure with the PSI monomer structure of the *Anabaena* PSI tetramer (PDB: 6JEO) clearly shows a structural correspondence between the C-terminal domain of IsiA2 and PsaL (Supplementary Fig. 5a–c), and the binding sites of Chls and Cars are conserved between the C-terminal domain of IsiA2 and PsaL (Supplementary Fig. 5c). The C-terminal domain of IsiA2 has a high sequence similarity to PsaL (Supplementary Fig. 5d); however, the amino acid residues of IsiA2-W426/F427/N451/W454 are remarkably different from the corresponding residues in PsaL (Supplementary Fig. 5e, f). These results provide evidence for the absence of PsaL in the PSI-IsiA structure, which may prevent the formation of PSI tetramer. Thus, PsaL is replaced by the C-terminal PsaL-like domain of IsiA2 in the *Anabaena* PSI-IsiA supercomplex.

The PSI core contains 92 Chls *a*, 21 *β*-carotenes, 3 [4Fe-4S] clusters, 2 phylloquinones, and 5 lipid molecules, which are summarized in Supplementary Table 3.

### Structure of the IsiA subunits

For the assignments of IsiA1-4, IsiA1-5, and IsiA2-1, we focused on the characteristic residues among the four types of IsiAs in *Anabaena.* IsiA1 is identified at positions 4 and 5 (Fig. 1a). The amino acid residues of F45/W47/K279/G281/V282/T283 in IsiA1 are significantly different from the corresponding residues in IsiA2, IsiA3, and IsiA5, allowing us to assign IsiA1-4 and IsiA1-5 (Supplementary Fig. 6a–c). IsiA1-5 binds 17 Chls *a* and 1 *β*-carotene (Fig. 2a). The axial ligands of the central Mg atom in a401, a402, a403, a405, a406, a408, a409, a410, a411, a412, a413, a414, and a415 of IsiA1-5 are coordinated by side chains of H207, H307, H96, H318, H221, H321, H31, Q34, N17, H145, H110, H144, and Q316, respectively (Fig. 2b). The Mg atom in a417 is associated with an oxygen atom of the carbonyl group of V282 (Fig. 2b). On the other hand, the ligands of a404, a407, and a416 seem to be water molecules which cannot be identified due to weak densities, and no amino acid residues existed in the vicinity of the central Mg atom of these Chls that are able to provide ligands to the Mg atoms.

**Fig. 2.**
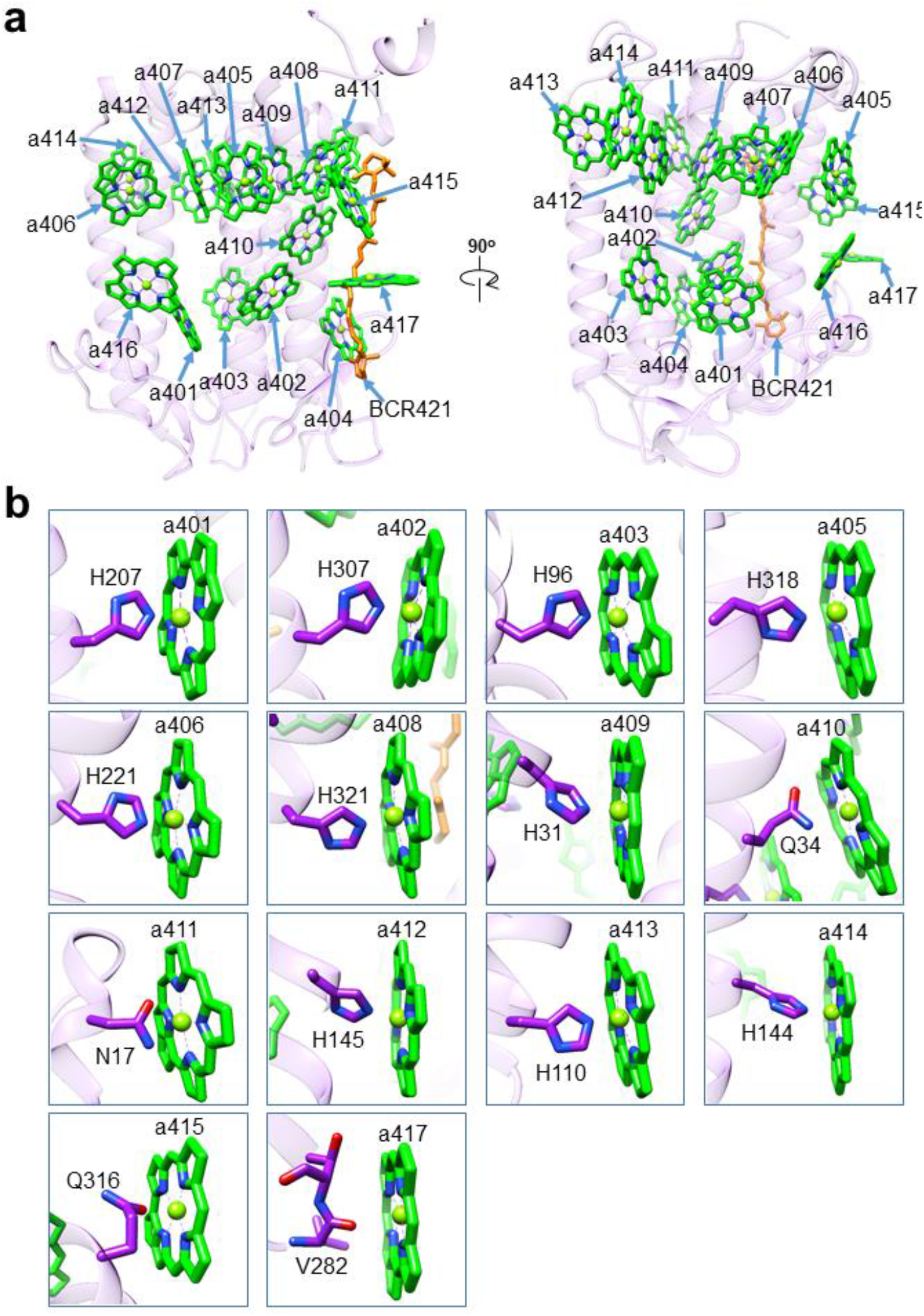
Structure of IsiA1-5. **a,** Structure of IsiA1-5 depicted in Cα atoms and arrangements of Chls and the *β*-carotene (BCR). **b,** Interactions of Chls with their ligands. Chls and the *β*-carotene are colored green and orange, respectively. Only rings of the Chl molecules are depicted.

IsiA1-4 binds only 10 Chls *a* and 1 *β*-carotene, which is largely different from that of IsiA1-5 (Supplementary Fig. 7a and Supplementary Table 3). The axial ligands of the central Mg atom in a401, a403, a408, a409, a411, a413, and a417 of IsiA1-4 are coordinated by the same amino acids as in IsiA1-5 (Supplementary Fig. 7b). On the other hand, the ligand of a404 seems to be a water molecule which cannot be identified due to weak densities, and no amino acid residues existed in the vicinity of the central Mg atom in the Chl molecule. The remarkable difference in the number of Chls between IsiA1-4 and IsiA1-5, albeit with the same gene product, may be due to weaker densities of IsiA1-4 compared with that of IsiA1-5 (Supplementary Fig. 3c; see Methods for the identification of pigment molecules), and/or the original absence of some Chls in IsiA1-4. This remarkable difference may bring consequences in excitation-energy transfer from IsiAs to the PSI core; however, a firm conclusion has to wait until a higher resolution structure is obtained.

The structure of IsiA2-1 can be separated into two parts, namely, an N-terminal IsiA domain and a C-terminal PsaL-like domain (Supplementary Fig. 5a–c). Amino-acid residues from the N-terminus to G341 are assigned to the N-terminal IsiA domain, because a global alignment of the *Anabaena* IsiA family shows that IsiA2-G341 corresponds to IsiA1-E354 and IsiA3-I352 which are the C-termini of the two latter genes (Supplementary Fig. 6a). On the other hand, IsiA2 has extra residues starting from N342 to the C-terminus (F476) (Supplementary Fig. 6a) which have some sequence similarities to the PsaL subunit of PSI, and thus are assigned to the C-terminal PsaL-like domain (Supplementary Fig. 5d).

The N-terminal IsiA domain of IsiA2 shows sequence similarity to IsiA1 (65%). The root mean square deviation (RMSD) between IsiA1-5 and IsiA2-1 is 0.95 Å for 302 Cα atoms (Supplementary Table 4). On the contrary, the C-terminal PsaL-like domain of IsiA2 exhibits sequence similarity to PsaL (62%) and structural similarity. The RMSD between PsaL and IsiA2-1 is 0.90 Å for 124 Cα atoms.

IsiA2-1 binds 17 Chls *a* and 5 *β*-carotenes (Fig. 3a, Supplementary Table 3). The axial ligands of the central Mg atom of a501, a502, a503, a505, a506, a508, a509, a510, a511, a512, a513, a531, and a532 are coordinated by side chains of H213, H300, H96, H311, H227, H314, H31, H34, N17, H145, H110, E375, and H380, respectively, whereas that of a533 is coordinated by a water molecule w25 (Fig. 3b). The Mg atom of a516 is associated with an oxygen atom of the carbonyl group of L191 (Fig. 3b). The ligands of a504 and a507 are not observed; they seem to be water molecules which cannot be identified due to weak densities. The binding properties of pigment molecules in the C-terminal PsaL-like domain of IsiA2-1 are virtually identical to those in PsaL (Supplementary Fig. 5c).

**Fig. 3.**
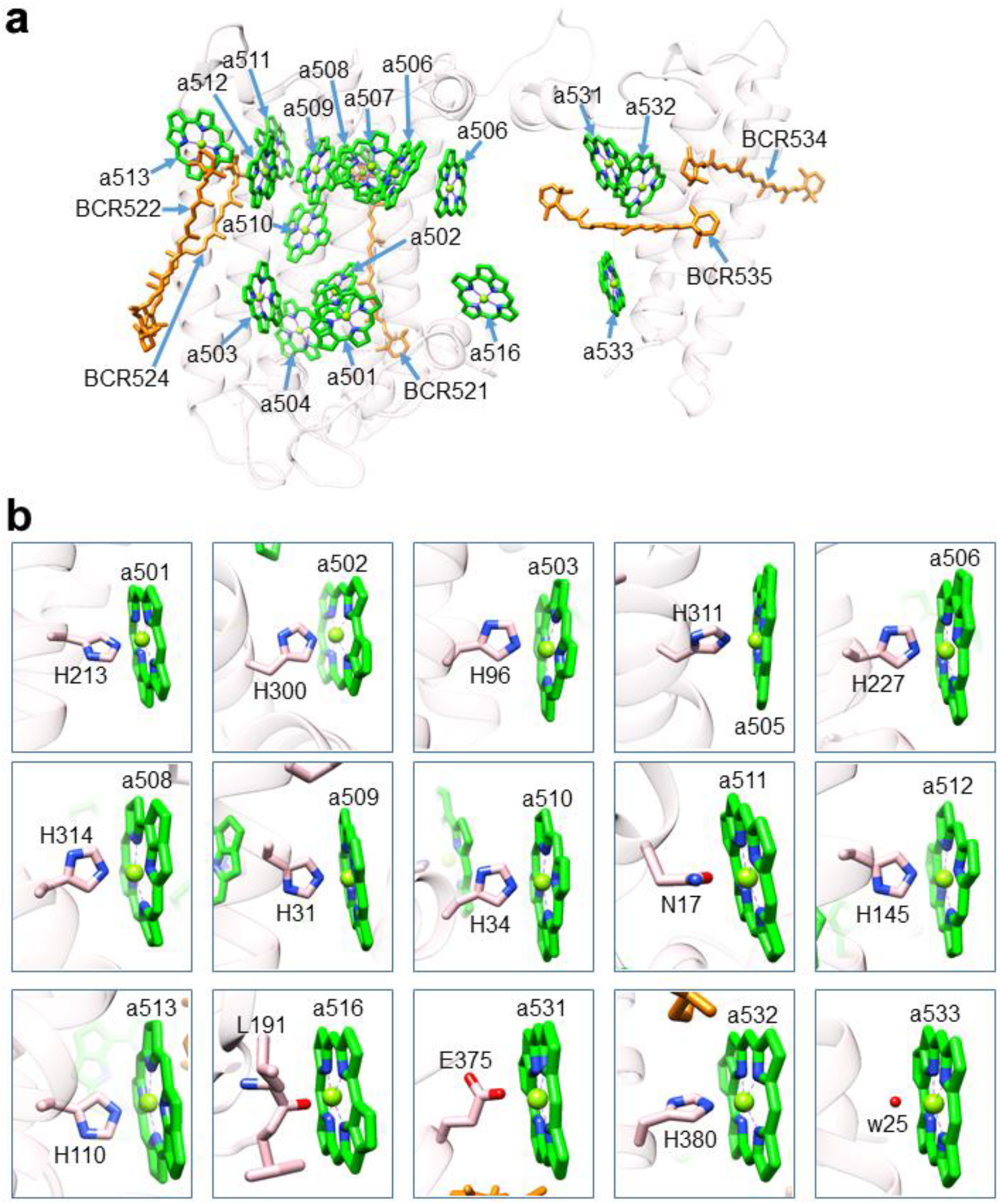
Structure of IsiA2-1. **a,** Structure of IsiA2-1 depicted in Cα atoms and arrangements of Chls and *β*-carotenes (BCRs). **b,** Interactions of Chls with their ligands. Chls and *β*-carotenes are colored green and orange, respectively. Only rings of the Chl molecules are depicted. w25, water molecule.

Among the unidentified IsiA subunits (IsiA-2, IsiA-3, and IsiA-6), IsiA-2 has 8 Chls *a*; IsiA-3 has 1 Chl *a* and 1 *β*-carotene; IsiA-6 has 11 Chls *a* (Supplementary Fig. 8, Supplementary Table 3). Each IsiA subunit shows different pigment compositions, which may be partly due to weak densities in these IsiA subunits (Supplementary Fig. 3c; see Methods for the identification of pigment molecules). The RMSD values between IsiA1-5 and IsiA-2/IsiA-3/IsiA1-4/IsiA-6 are 0.56–0.95 Å for 277–326 Cα atoms (Supplementary Table 4).

### Interactions among the IsiA subunits

There are many interactions among the IsiA subunits, which are mostly hydrophobic (Fig. 4a). Amino-acid residues A101/A105/L108 and BCR424 in IsiA1-4 interact with Chl molecules of a405/a415 in IsiA1-5 through hydrophobic interactions at distances of 3.3–3.9 Å (the upper panel in Fig. 4b). The amino-acid residues A44/F45/W47/F48/S51 and the pigment molecules a404 and BCR424 in IsiA1-4 are also associated with F254/Y258/L278/K279/F280 and a417 in IsiA1-5 through hydrophobic interactions at distances of 3.3–4.7 Å (the lower panel in Fig. 4b). The pigment molecule BCR424 in IsiA-3 interacts with L251/F254/V255/Y258 in IsiA1-4 through hydrophobic interactions at distances of 3.5–4.2 Å (Fig. 4c). The amino-acid residues L40/A44/F45/A105/L108 in IsiA1-5 are associated with a405/a415/a417 in IsiA-6 through hydrophobic interactions at distances of 3.1-3.8 Å (Fig. 4d). The amino-acid residues L40/A44/A105/I108/L113 and BCR524 in IsiA2-1 are coupled with a405/a415/a417 in IsiA-2 through hydrophobic interactions at distances of 3.4–4.0 Å (Fig. 4e). No characteristic interactions between IsiA-2 and IsiA-3 are found in the present structure (black square in Fig. 4a). It should be noted that other interactions remain ambiguous because of weak densities in the corresponding map among IsiAs (Supplementary Fig. 3c).

**Fig. 4.**
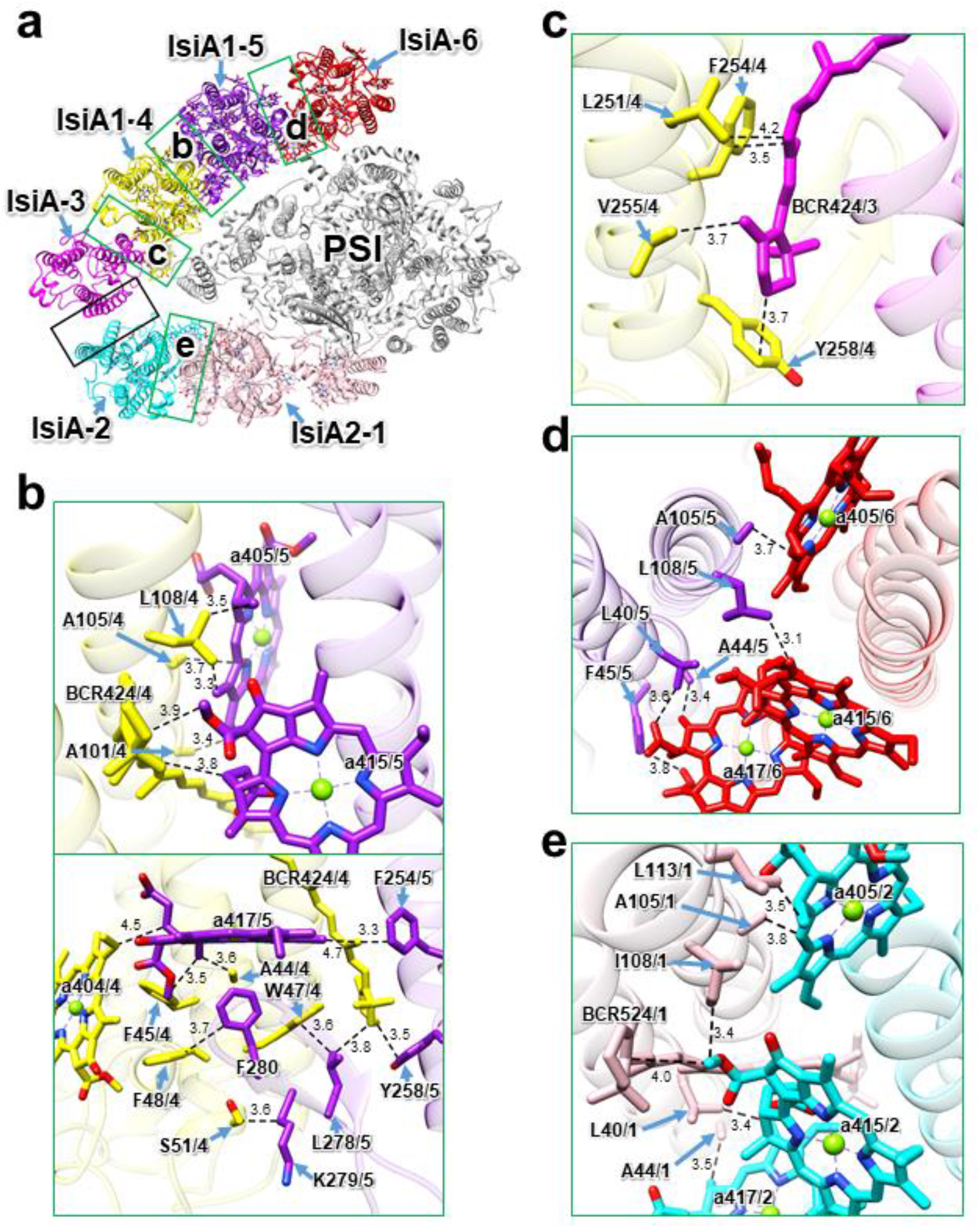
Interactions among the IsiA subunits. **a,** The structure of PSI-IsiA viewed from the cytosolic side. The green square areas are enlarged in panels **b**–**e**, whereas the black squared area does not have a characteristic interaction in the present structure. **b**, Interactions between IsiA1-4 and IsiA1-5 (upper and lower panels). **c**, Interactions between IsiA-3 and IsiA1-4. **d**, Interactions between IsiA1-5 and IsiA-6. **e**, Interactions between IsiA2-1 and IsiA-2. Interactions are indicated by dashed lines, and the numbers are distances in Å. The letters represent the numbering of amino acid residues and pigments in each IsiA subunit; for example, A105/4 means Ala105 in IsiA1-4; a415/5 means Chl *a* 415 in IsiA1-5. BCR, *β*-carotene.

### Interactions between the IsiA subunits and the PSI core

The interactions between IsiAs and PSI are shown in Fig. 5. The amino-acid residues W313/V331/L347 in IsiA2-1 interact with F333 and the Chl molecules a833/a846 in PsaA through hydrophobic interactions at distances of 3.4–3.7 Å, and an oxygen atom of the carbonyl group of G341 in IsiA2-1 is hydrogen-bonded with a nitrogen atom of T16 in PsaD at a distance of 3.3 Å (the left panel of Fig. 5b). Moreover, BCR521 in IsiA2-1 is associated with a837 in PsaA at a distance of 3.6 Å (the right panel of Fig. 5b). The amino-acid residue I333 in IsiA1-4 interacts with BCR858 in PsaA at a distance of 4.4 Å (Fig. 5c), whereas W14 in IsiA1-4 is located near a821 in PsaA at a distance of 3.5 Å (Fig. 5d). It should be noted that other interactions remain ambiguous because of weak densities in the corresponding map between IsiAs and PSI (Supplementary Fig. 3c).

**Fig. 5.**
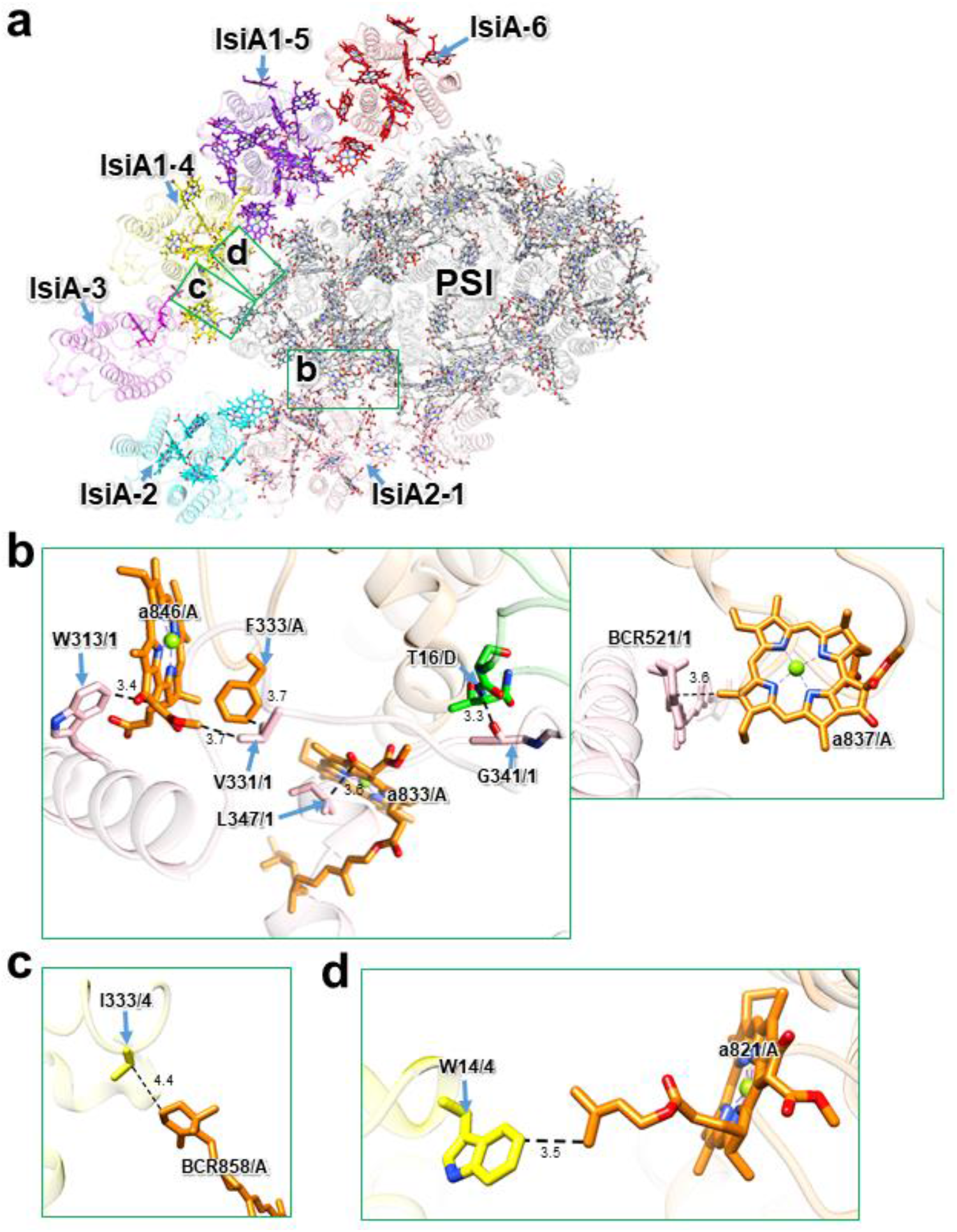
Interactions between IsiAs and PSI. **a,** The structure of PSI-IsiA viewed from the cytosolic side. Green squared areas are enlarged in panels **b–d**. **b**, Interactions between IsiA2-1 and PsaA/PsaD (left and right panels). **c**, Interaction between residue I333 of IsiA1-4 and BCR858 of PsaA. **d**, Interaction between residue W14 of IsiA1-4 and Chl a821 of PsaA. Interactions are indicated by dashed lines, and the numbers are distances in Å. The letters represent the numbering of amino acid residues and pigments in each subunit of IsiAs/PsaA/PsaD; for example, W313/1 means Typ313 in IsiA2-1; a846/A means Chl *a* 846 in PsaA. BCR, *β-* carotene.

### Excitation-energy-transfer processes in the PSI-IsiA supercomplex

Time-resolved fluorescence (TRF) spectra of the PSI-IsiA supercomplex were measured at 77 K and globally analyzed to obtain fluorescence decay-associated (FDA) spectra (Fig. 6). The 55-ps FDA spectrum exhibits positive amplitudes around 685 nm and negative amplitudes around 727 nm. Since a set of positive and negative bands indicates energy transfer from Chl with the positive one to Chl with the negative one, the positive-negative pair of the 685 and 727-nm bands reflects energy transfer from the 685-nm component to the 727-nm component. The 55-ps FDA spectrum also shows a positive shoulder around 694 nm and a negative shoulder around 707 nm. The 120-ps FDA spectrum displays two positive bands at 690 and 707 nm. This is in striking contrast to the previous results of the *Anabaena* PSI monomer, dimer, and tetramer, in which only a broad band around 728 nm appears in the 100–170-ps FDA spectra^32^. These results suggest that the *Anabaena* IsiAs affected the excitation-energy-transfer processes occurring in the early time region after excitation.

**Fig. 6.**
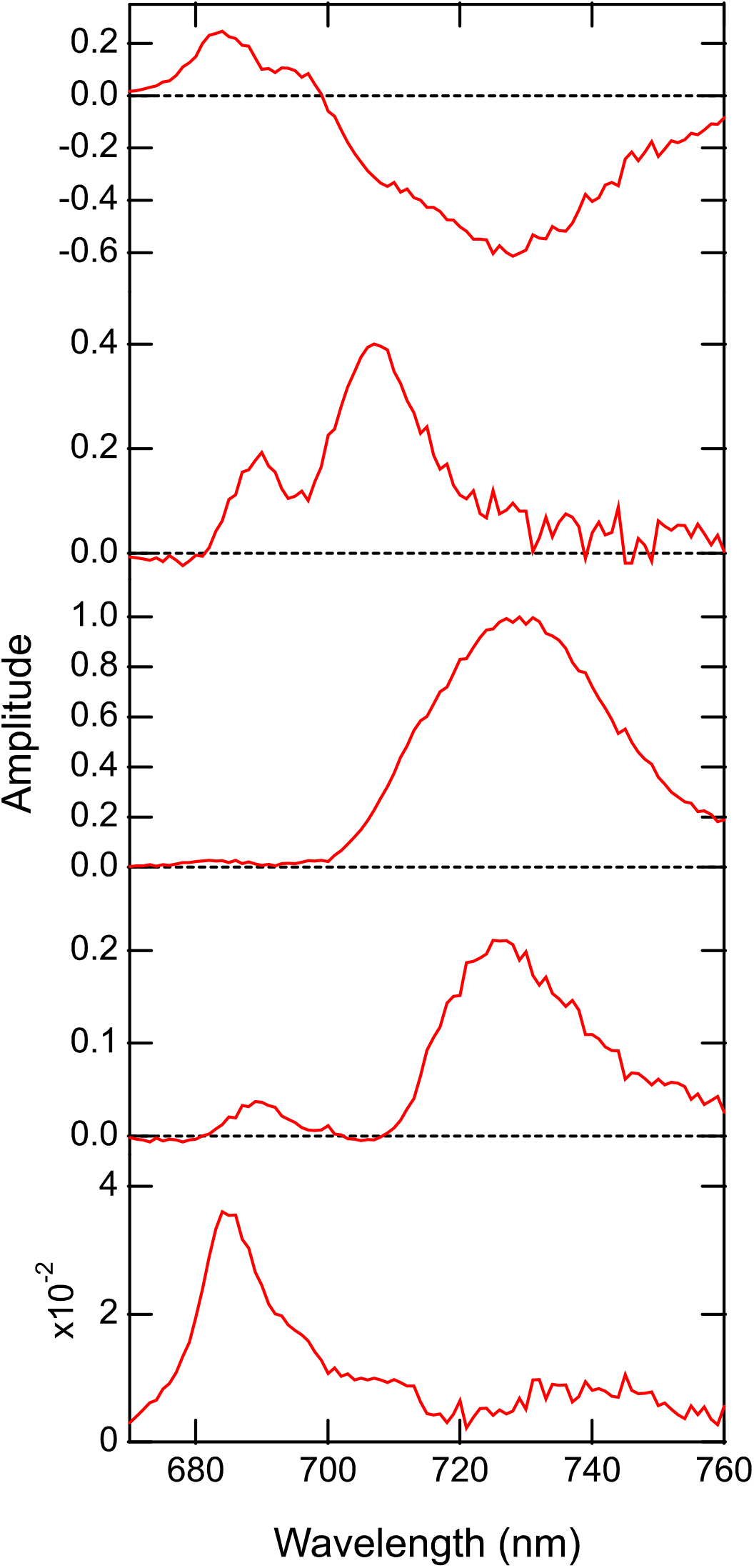
Fluorescence decay-associated (FDA) spectra of PSI-IsiA. Time-resolved fluorescence (TRF) spectra were measured at 77 K with excitation at 445 nm, followed by global analysis to construct the FDA spectra. The analyzed time constants for the spectra are 55 ps, 120 ps, 520 ps, 1.2 ns, and 3.9 ns from top to bottom. The FDA spectra were normalized by the maximum amplitude of the 520-ps spectrum.

## Discussion

### Multiple expressions of the *Anabaena* IsiA family and its structures

Our previous study of the two-dimensional BN/SDS-PAGE analysis using thylakoids from the wild-type strain of *Anabaena* grown under the iron-deficient condition showed the detection of only IsiA1 subunit in the fraction of the PSI-IsiA supercomplex^30^. However, the present study showed the expression of IsiA2, IsiA3, and IsiA5 as well as IsiA1 at transcript and protein levels (Supplementary Fig. 1a, c). However, the band density of IsiA1 on the SDS-PAGE gel using the purified PSI-IsiA supercomplexes is remarkably higher than those of IsiA2, IsiA3, and IsiA5 (Supplementary Fig. 1c). This may be one of the reasons for the failure of detection of IsiA2, IsiA3, and IsiA5 in our previous analysis^30^. The expression of all *isiA* genes in *Anabaena* is in good agreement with the results of *Leptolyngbya* showing that the five types of IsiA proteins were biosynthesized under an iron-deficient condition^29^. Thus, cyanobacteria with more than one *isiA* gene may accumulate all *isiA* products under iron-deficient conditions.

The structure of the PSI-IsiA supercomplex reveals the binding of six IsiA subunits to the PSI-monomer core, three of which were identified as IsiA2-1, IsiA1-4, and IsiA1-5, whereas the remaining three subunits (IsiA-2, IsiA-3, and IsiA-6) were not identified yet (Fig. 1a). The structures of the five IsiA subunits other than IsiA2-1 are similar to that of the structurally known IsiA subunit of other cyanobacteria, with the characteristic six trans-membrane helices^23–25^. This is in good agreement with the result of the sequence alignment that puts the *Anabaena* IsiA1 into the same group of the structurally known IsiA family^30^. The *Anabaena* IsiA1 has also a high sequence similarity with the *Anabaena* IsiA2, IsiA3, and IsiA5 (Supplementary Fig. 6a), with RMSDs between IsiA1-5 and IsiA2-1/IsiA-2/IsiA-3/IsiA1-4/IsiA-6 in the range of 0.56–0.95 Å (Supplementary Table 4). Because all four IsiAs in *Anabaena* are expressed under our experimental conditions (Supplementary Fig. 1a, c), these findings indicate that some of the unidentified subunits of IsiA-2, IsiA-3, and IsiA-6 are at least the gene products of *isiA3* and *isiA5* in the PSI-IsiA supercomplex.

### Competitive incorporation of IsiA2 or PsaL into PSI under iron-deficient conditions

Among the six IsiA subunits, IsiA2-1 shows an unusual structure different from the other five IsiA subunits, because of its C-terminal PsaL-like domain (Fig. 3a, Supplementary Fig. 5). Shen and co-workers showed that *Leptolyngbya* had IsiA4 containing a PsaL-like domain^29^, similar to the *Anabaena* IsiA2^30^. The authors also suggested that the *Leptolyngbya* IsiA4 was related to protein aggregation and the formation of supercomplexes with PSI^29^. These observations indicate that cyanobacteria having IsiAs containing a PsaL-like domain organize PSI-monomer-IsiA supercomplexes through the replacement of PsaL with the PsaL-containing IsiAs under iron-deficient conditions.

It is interesting to note that the *Anabaena* PsaL is still expressed under the iron-deficient condition and then contributes to the formation of PSI tetramers. This is because the PSI tetramers were detected by two-dimensional BN/SDS-PAGE using the *Anabaena* thylakoids under the iron-deficient condition^30^. Since PsaL plays a crucial role in the oligomerization of PSI^33–35^, it seems that PsaL competes with IsiA2 for interactions with PSI and for the formation of PSI monomer or PSI tetramer in *Anabaena* under iron-deficient conditions.

### Molecular assembly of the PSI-monomer-IsiA supercomplex

Based on these observations, we propose a schematic model for the assembly of the PSI-monomer-IsiA supercomplex and PSI tetramer in *Anabaena* under iron-deficient conditions (Fig. 7). Two subunits of PsaL and IsiA2 play an important role in determining the oligomeric states of PSI in *Anabaena.* When IsiA2 is expressed and bound to PSI monomer without PsaL, oligomerization of monomeric PSI to dimers and tetramers may be inhibited due to the inability of PsaL to bind to the PSI monomer. After the binding of IsiA2 at position 1, the PSI-IsiA2-1 supercomplex assembles into the PSI-monomer-IsiA supercomplex with the association of the remaining five IsiA subunits. As a result, no PSI-tetramer-IsiA fraction was obtained by the two-dimensional BN/SDS-PAGE^30^. In contrast, when PsaL is first bound to PSI monomer without PsaL and IsiAs, further assembly to PSI tetramers may proceed irrespective of the expression of IsiA2. The competitive assembly of oligomeric PSI cores and a PSI-monomer-IsiA supercomplex would often occur in cyanobacteria expressing both PsaL and IsiAs in iron-limited environments.

**Fig. 7.**
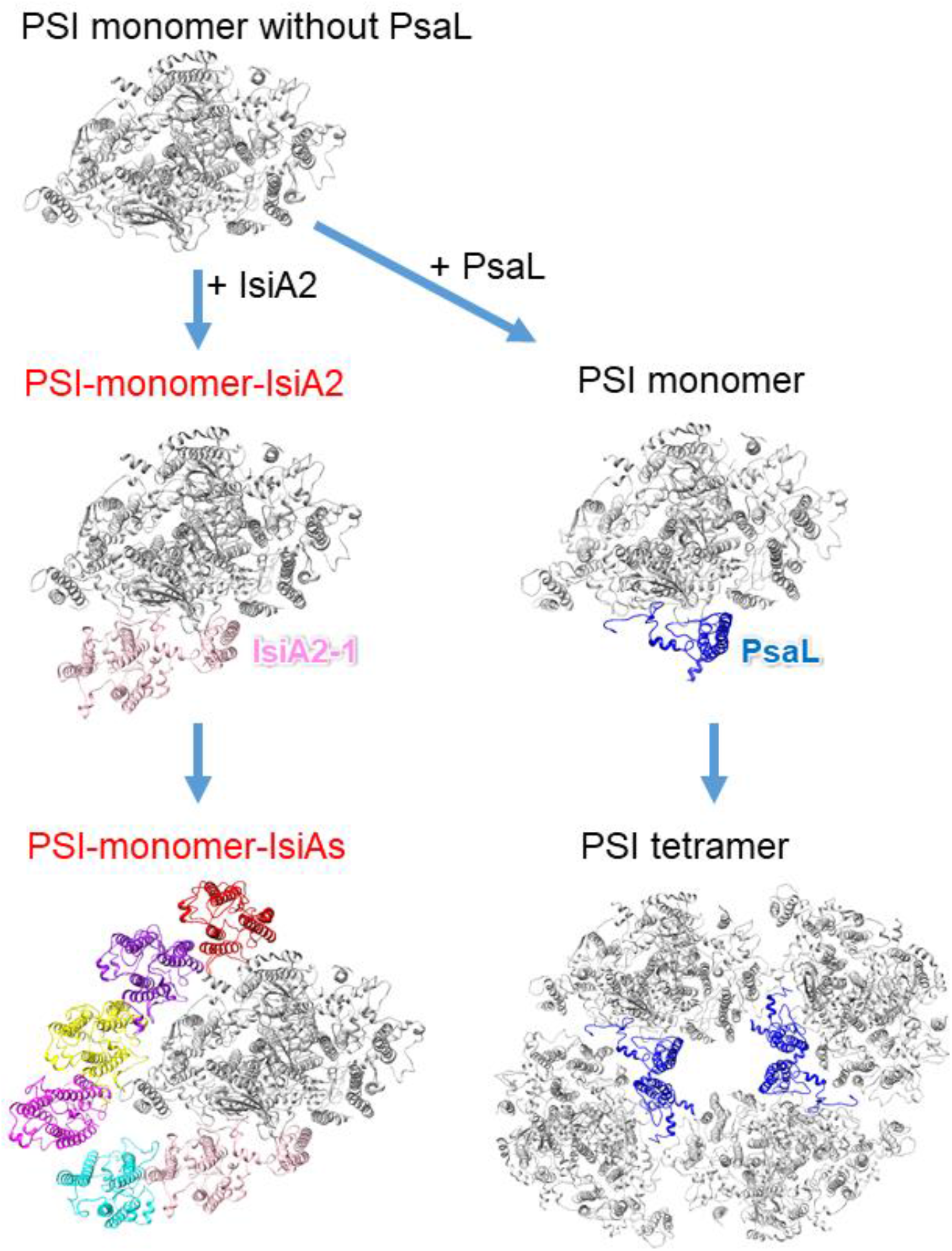
An assembly model proposed for the *Anabaena* PSI-IsiA supercomplex. The structures are viewed from the cytosolic side. The PSI monomer and tetramer are taken from 6JEO (PDB ID).

### Functional roles of IsiAs in excitation-energy transfer

The binding of IsiAs to PSI affects the absorption and fluorescence spectra (Supplementary Fig. 1e, f). The absorption spectrum of PSI-IsiA exhibits the Qy band of Chl *a* at 673 nm (Supplementary Fig. 1e), which is shorter than that in the spectrum of the PSI monomer from *Anabaena*^11^. This is characteristic of the existence of IsiAs, as observed in the PSI-IsiA supercomplexes from various cyanobacteria^21,26,28^. Such a blue-shifted band has also been observed in the absorption spectrum of thylakoid membranes prepared from this cyanobacterium grown under the iron-deficient condition^30^. Furthermore, the fluorescence-emission spectrum of PSI-IsiA shows two bands at 687 and 727 nm (Supplementary Fig. 1f). The former appears to originate from IsiAs as observed in the *Anabaena* thylakoids^30^ and other cyanobacteria^21,36,37^, whereas the latter is one of the specific low-energy Chls in PSI, which have been denoted as Low2 based on its absence in *Gloeobacter violaceus* PCC 7421 and *Synechocystis* sp. PCC 6803^38^. In contrast, the fluorescence spectrum of the PSI monomer of *Anabaena* grown under iron-replete conditions has shown a band at 730 nm, without the characteristic 687-nm band observed in the PSI-IsiA spectrum^11,32^. Thus, the spectroscopic properties characteristic of the *Anabaena* IsiAs clearly appear even under steady-state experimental conditions.

The excitation-energy dynamics in the *Anabaena* PSI-IsiA supercomplex have been examined by the FDA spectra (Fig. 6). The characteristic fluorescence band at 684 nm originating from IsiAs decays with a time constant of 55 ps, and the corresponding rise of fluorescence appears around 727 nm, suggesting excitation-energy transfer from IsiAs to PSI. In addition to the negative 727-nm band, a negative shoulder is recognized around 707 nm in the 55-ps FDA spectrum, which is followed by a positive 707-nm band in the 120-ps FDA spectrum. Since the distinct positive band at 707 nm was not observed in the PSI monomer^39^, it is suggested that the 707-nm band occurs in the interface between IsiAs and PSI. In contrast to the 55-ps FDA spectrum exhibiting the positive-negative pair, the 690-nm band lacking a corresponding negative band in the 120-ps FDA spectrum may be attributed to excitation-energy quenching through interactions among pigments around/within IsiAs. Energy quenching with time constants of hundreds of picoseconds has been interpreted by various spectroscopic studies using photosynthetic pigments and LHCs^40,41^. Thus, the *Anabaena* PSI-IsiA may possess quenching sites at 690 and 707 nm, the latter having the same transition energy as found in the *Anabaena* PSI tetramer^11,32^. The 686-nm band appears in the 3.9-ns FDA spectrum, suggesting that uncoupled Chls in IsiAs cannot transfer excitation energy to other pigments, which is similar to that observed in the *T. vulcanus* PSI-IsiA^25^.

Based on the properties of excitation-energy-transfer processes and the structure of the *Anabaena* PSI-IsiA, we propose excitation-energy-transfer pathways from IsiAs to PSI (Supplementary Fig. 9). The excitation energy in IsiA2-1, IsiA1-4, and IsiA-6 appear to be directly transferred to the PSI core, due to the close pigment-pigment interactions of these IsiA subunits with PSI subunits (Supplementary Fig. 9a). On the contrary, the energy in IsiA-2, IsiA-3, and IsiA-5 may be transferred once to the neighboring IsiA subunits prior to excitation-energy transfer to PSI. As excitation-energy transfer from IsiAs to PSI occur with a time constant of 55 ps (Fig. 6), Chl couplings of a516/a533 within IsiA2-1, IsiA2-1-a508/PsaA-a846, IsiA1-4-a415/PsaA-a845, IsiA1-4-a417/PsaA-a845, IsiA1-4-a404/PsaA-a816, and IsiA-6-a417/PsaK-a101 (Supplementary Fig. 9b–f) may serve as energy donors and acceptors between IsiAs and PSI. As for IsiA2-1, the pigment molecules in the C-terminal PsaL-like domain may function similarly to those in PsaL, because of the almost same pigment arrangements between the C-terminal domain of IsiA2-1 and PsaL (Supplementary Fig. 5b, c). Therefore, the Chl couplings between the N-terminal domain and C-terminal domains within IsiA2-1 may be involved in excitation-energy transfer with a time constant of 55 ps. Furthermore, the Car-Chl coupling of IsiA2-1-BCR521/PsaA-a837 (Supplementary Fig. 9c) may contribute to ultrafast excitation-energy transfer in the time order of femtoseconds as observed in ultrafast spectroscopies^40^ and energy quenching by Chl-Car interactions with a time constant of 120 ps, between IsiA2-1 and PsaA.

### Structural comparisons of IsiAs between *Anabaena* and other cyanobacteria

In the PSI-trimer-IsiA structures, six IsiA subunits were associated with the PSI-monomer unit^23–25^. Here, we compare the binding properties of IsiAs between *Anabaena* and other cyanobacteria (Supplementary Fig. 10). The PSI-monomer cores are well fitted between *Anabaena* and *Synechocystis* sp. PCC 6803 (Supplementary Fig. 10a). The six IsiA subunits in the *Synechocystis* PSI-IsiA are temporally named IsiA-1 to IsiA-6 (Supplementary Fig. 10a). The IsiA1-4, IsiA1-5, and IsiA-6 subunits in the *Anabaena* PSI-IsiA are located at similar positions as the IsiA-1, IsiA-2, and IsiA-3 subunits, respectively, in the *Synechocystis* PSI-IsiA; these three IsiAs are associated with PsaA in the two species (Supplementary Fig. 10a). However, the remaining three IsiA subunits are bound to the outside of PsaB in the *Synechocystis* PSI-IsiA, whereas they are bound to the remaining part of PsaA opposite to the side of PsaB in the *Anabaena* PSI-IsiA. This may be due to differences in the structures of the IsiA1-4, IsiA1-5 and IsiA-6 subunits in the *Anabaena* PSI-IsiA with the corresponding IsiA subunits in the *Synechocystis* PSI-IsiA (Supplementary Fig. 10b). In particular, the structural gap of IsiA-6 is much larger than that of IsiA1-4 and IsiA1-5 (Supplementary Fig. 10b). The structural distortion of the *Anabaena* IsiAs may also occur by the unusual binding of IsiA2-1, IsiA-2, and IsiA-3 to PSI. Moreover, the C-terminal loop of IsiA is missing in the IsiA1-4, IsiA1-5, and IsiA-6 subunits in the *Anabaena* PSI-IsiA (yellow arrows in Supplementary Fig. 10b). These structural differences are observed in the structural comparisons of PSI-IsiA between *Anabaena* and *Thermosynechococcus vulcanus* NIES-2134 (Supplementary Fig. 10c) and between *Anabaena* and *Synechococcus elongatus* PCC 7942 (Supplementary Fig. 10d), which may lead to a suppression of the binding of IsiAs to the outside of PsaB in *Anabaena*.

## Conclusions

This study demonstrates the overall structure of a PSI-monomer-IsiA supercomplex from *Anabaena* grown under the iron-deficient condition. The structure of IsiA2-1 shows both a typical N-terminal IsiA domain and a C-terminal PsaL-like domain, and is bound to PSI by substituting PsaL with the C-terminal PsaL-like domain of IsiA2. The binding of IsiA2 to PSI leads to inhibition of PSI oligomerization, resulting in a monomeric PSI core with six IsiA subunits bound. Unlike the typical PSI-trimer-IsiA structures, the IsiA subunits are associated with the PsaA side but not with the outside of PsaB in the *Anabaena* PSI-IsiA structure. This may be attributed to the differences in interactions among IsiAs between *Anabaena* and other cyanobacteria having the PSI-trimer-IsiA structures. These structural findings may be characteristic of cyanobacteria having multiple copies of *isiA* genes, and hence, may contribute to a survival strategy for such types of cyanobacteria including *Anabaena* under iron-limited conditions.

## Acknowledgements

We thank Ms. Kumiyo Kato for her helpful assistance in this study. This work was supported by JSPS KAKENHI grant Nos. JP20H02914 (Koji.K.), JP21K19085 (R.N.), JP20K06528 (Keisuke.K), and JP17H06434 and JP22H04916 (J.-R.S.), JST-Mirai Program Grant Number JPMJMI20G5 (K.Y.), Takeda Science Foundation (Koji.K.), and the Cyclic Innovation for Clinical Empowerment (CiCLE) from the Japan Agency for Medical Research and Development, AMED (T.H., Keisuke.K., K.Y.).

## Author Contributions

R.N. conceived the project; R.N., N.T., and S.S. prepared the PSI-IsiA supercomplex and analyzed its biochemical characterization; S.E. performed transcription analysis of IsiAs; Y.U., M.F., and S.A. performed TRF measurements of the PSI-IsiA supercomplex and their data analysis; T.S. and N.D. identified subunits in the PSI-IsiA supercomplex; T.H. collected cryo-EM images; Koji.K. processed the cryo-EM data and reconstructed the final cryo-EM map; Koji.K. built the structural model and refined the final model; Y.N. analyzed structural data; Keisuke.K. commented on the structural data; K.Y. and J.-R.S. supervised this project; R.N. and S.A. wrote the draft manuscript; and R.N. and J.-R.S. wrote the final manuscript, and all of the authors joined the discussion of the results.

## Methods

### Cell growth and preparation of thylakoid membranes

The *Anabaena* cells were grown in an iron-replete BG11 medium supplemented with 10 mM HEPES-KOH (pH 8.0) at a photosynthetic photon flux density of 30 μmol photons m^-2^ s^-1^ at 30°C with bubbling of air containing 3% (v/v) CO_2_^30^. For iron starvation, the cells grown under the iron-replete condition were substituted with an iron-free BG11 medium and then cultured according to the method of Nagao et al.^30^. The cells were harvested by centrifugation, and then thylakoid membranes were prepared by agitation with glass beads^39^, followed by suspension with a 20 mM MES-NaOH (pH 6.5) buffer containing 0.2 M trehalose, 5 mM CaCl_2_, and 10 mM MgCl_2_ (buffer A).

### Transcription analysis of IsiAs

The cells were grown for 0 and 20 days under the iron-deficient condition, and were collected by centrifugation at 4 °C and stored at −80 °C until use. Total RNA was extracted from the cells with a PGTX solution^42^ and purified using NucleoSpin RNA (MACHEREY-NAGEL). cDNA was synthesized from 1 μg of total RNA using ReverTra Ace qPCR RT Master Mix (TOYOBO), including random and oligo (dT) primers. The presence of contaminating genome DNA was confirmed by incubating 1 μg of total RNA without reverse transcriptase under the condition of cDNA synthesis. According to the manufacturer’s instructions, PCR was conducted with cDNA as the template using Tks Gflex DNA Polymerase (Takara Bio). Quantitative reverse transcription PCR (qRT-PCR) was performed according to the method of Ehira and Miyazaki^43^ using Go Taq qPCR Master Mix (Promega). Primers used for PCR and qRT-PCR are listed in Supplementary Table 5.

### Purification of the PSI-IsiA supercomplex

Thylakoid membranes were solubilized with 1% (w/v) *n*-dodecyl-*β*-D-maltoside *(β-* DDM) at a Chl concentration of 0.25 mg mL^-1^ for 10 min on ice in the dark with gentle stirring. After centrifugation at 100,000 × g for 10 min at 4°C, the resultant supernatant was loaded onto a Q-Sepharose anion-exchange column (1.6 cm of inner diameter and 25 cm of length) equilibrated with buffer A containing 0.03% *β*-DDM (buffer B). The column was washed with buffer B containing 50 mM NaCl until the eluate became colorless. Two types of buffers, buffer B and buffer C (buffer B containing 500 mM NaCl), were used for the elution of PSI-IsiA from the column in a linear-gradient step of 0–600 min, 10–50% buffer C. The flow rate was set to 2.0 mL min^-1^. The PSI-IsiA-enriched fraction was eluted at around 200–230 mM NaCl.

The eluted PSI-IsiA were precipitated by centrifugation after the addition of polyethylene glycol 1500 to a final concentration of 15% (w/v), and then suspended with Buffer B. The resultant PSI-IsiA samples were loaded onto a linear gradient containing 10–40% (w/v) trehalose in a medium of 20 mM MES-NaOH (pH 6.5), 5 mM CaCl_2_, 10 mM MgCl_2_, 100 mM NaCl, and 0.03% *β*-DDM. After centrifugation at 154,000 × g for 18 h at 4°C (P40ST rotor; Hitachi), a major green fraction (Supplementary Fig. 1b) was collected and concentrated using a 150 kDa cut-off filter (Apollo; Orbital Biosciences) at 4,000 × g. The concentrated samples were stored in liquid nitrogen until use.

### Biochemical and spectroscopic analyses of the PSI-IsiA supercomplex

Subunit composition of the PSI-IsiA supercomplex was analyzed by SDS-polyacrylamide gel electrophoresis (PAGE) containing 16% (w/v) acrylamide and 7.5 M urea according to the method of Ikeuchi and Inoue^44^ (Supplementary Fig. 1c). The PSI-IsiA supercomplexes (4 μg of Chl) were solubilized by 3% lithium lauryl sulfate and 75 mM dithiothreitol for 10 min at 60°C, and loaded onto the gel. A standard molecular weight marker (SP-0110; APRO Science) was used. The subunit bands were assigned by mass spectrometry according to the method of Nagao et al.^45^. Oligomeric state of the PSI-IsiA supercomplex was analyzed by BN-PAGE containing 3–12% acrylamide according to the method of Schägger and von Jagow^46^ (Supplementary Fig. 1d), with 2.5 μg of Chl loaded onto the gel. As a reference, thylakoid membranes prepared from this cyanobacterium under the iron-replete condition were solubilized with 1% *β*-DDM at a Chl concentration of 0.25 mg mL^-1^ for 10 min on ice in the dark. After centrifugation at 20,000 *×* g for 10 min at 4°C, the resultant supernatant (2.5 μg of Chl) was loaded onto the gel. Pigment compositions of the PSI-IsiA supercomplex were analyzed by HPLC according to the method of Nagao et al.^47^, and the elution profile was monitored at 440 nm (Supplementary Fig. 1g).

Absorption spectrum was measured at 298 K using a UV-Vis spectrophotometer (UV-2450; Shimadzu) (Supplementary Fig. 1e). Steady-state fluorescence-emission spectrum was measured at 77 K using a spectrofluorometer (RF-5300PC; Shimadzu) (Supplementary Fig. 1f). TRF was recorded by a time-correlated single-photon counting system with a wavelength interval of 1 nm and a time interval of 2.44 ps^48^. A picosecond pulse diode laser (PiL044X; Advanced Laser Diode Systems) was used as an excitation source, and it was operated at 445 nm with a repetition rate of 3 MHz. The TRF-measurement conditions were described in detail^49^. The fluorescence intensities were obtained as a function of time (*t*) and wavelength (*λ*), and globally analyzed with time constants (*τ*) independent of *λ*, as 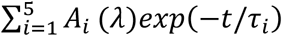. Here, *A_i_*(*λ*) is the FDA spectrum with *τ_i_*. The weighted residual map between the measured and calculated data is shown in Supplementary Fig. 11.

### Cryo-EM data collection

For cryo-EM experiments, 3-μL aliquots of the *Anabaena* PSI-IsiA supercomplex (0.53 mg Chl mL^-1^) in a 20 mM MES-NaOH (pH 6.5) buffer containing 5 mM CaCl_2_, 10 mM MgCl_2_, 100 mM NaCl, and 0.03% *β*-DDM were applied to Quantifoil R0.6/1, Cu 200 mesh grids pre-treated by gold sputtering. Without waiting for incubation, the excess amount of the solution was blotted off for 6 sec with a filter paper in an FEI Vitrobot Mark IV at 4°C under 100% humidity. The grids were plunged into liquid ethane cooled by liquid nitrogen and then transferred into a CRYO ARM 300 electron microscope (JEOL) equipped with a cold-field emission gun operated at 300 kV. AI detection of hole positions was carried out with yoneoLocr^50^, which prevented the stage alignment failure. All image stacks acquired were collected from 5 × 5 holes per stage adjustment to the central hole using SerialEM^51^ and JAFIS Tool version 1 (developed by Dr. Bartosz Marzec, JEOL), which synchronized image shifts with beam tilts, objective stigmas for removal of axial coma aberrations, and two-fold astigmatism. The images were zero-loss energy filtered and recorded at a nominal magnification of × 100,000 on a direct electron detection camera (Gatan K3, AMETEK) with a nominal defocus range of −1.8 to −0.8 μm. One-pixel size corresponded to 0.495 Å. Each image stack was exposed at a dose rate of 13.6 e^-^Å^-2^sec^-1^ for 3.0 sec in CDS mode, and consisted of dose-fractionated 50 movie frames. In total 22,575 image stacks were collected.

### Cryo-EM image processing

The resultant movie frames were aligned and summed using MotionCor2^52^ to yield dose weighted images. Estimation of the contrast transfer function (CTF) was performed using CTFFIND4^53^. All of the following processes were performed using RELION3.1^54^. In total 1,775,806 particles were automatically picked up and used for reference-free 2D classification. Then, 280,489 particles were selected from good 2D classes and subsequently subjected to 3D classification without imposing any symmetry. An initial model for the first 3D classification was generated *de novo* from 2D classifications. As shown in Supplementary Fig. 2c, the final PSI-IsiA structure was reconstructed from 47,602 particles. The overall resolution of the cryo-EM map was estimated to be 2.62 Å by the gold-standard FSC curve with a cut-off value of 0.143 (Supplementary Fig. 3a)^55^. Local resolutions were calculated using RELION (Supplementary Fig. 3c).

### Model building and refinement

Two types of the cryo-EM maps were used for the model building of the PSI-IsiA supercomplex: one is a postprocessed map, and the other is a denoised map using Topaz version 0.2.4^56^. The postprocessed map was denoised using the trained model in 100 epochs with two half-maps. In particular, the three subunits of IsiA-2, IsiA-3, and IsiA-6 were modeled with polyalanines using the denoised map (Supplementary Fig. 12). Each subunit of the homology models constructed using the Phyre2 server^57^ was first manually fitted into the two maps using UCSF Chimera^58^, and then their structures were inspected and manually adjusted against the maps with Coot^59^. Each model was built based on interpretable features from the density maps with the contour levels of 1.0 and 2.5 σ in the denoised and postprocessed maps, respectively. The complete PSI-IsiA structure was refined with phenix.real_space_refine^60^ and Servalcat^61^ with geometric restraints for the protein-cofactor coordination. The final model was validated with MolProbity^62^, EMRinger^63^, and *Q*-score^64^. The statistics for all data collection and structure refinement are summarized in Supplementary Table 1, 2. All structural figures were made by PyMOL^65^ and UCSF Chimera.

### Data availability

Atomic coordinate and cryo-EM maps for the reported structure of the PSI-IsiA supercomplex have been deposited in the Protein Data Bank under an accession code 7Y3F and in the Electron Microscopy Data Bank under an accession code EMD-33593.

## References

1 Blankenship, R. E. Molecular Mechanisms of Photosynthesis. 3rd edn, (Wiley-Blackwell, 2021).

2 Umena, Y., Kawakami, K., Shen, J.-R. & Kamiya, N. Crystal structure of oxygen-evolving photosystem II at a resolution of 1.9 Å. Nature 473, 55–60 (2011).

3 Shen, J.-R. The structure of photosystem II and the mechanism of water oxidation in photosynthesis. Annu. Rev. Plant Biol. 66, 23–48 (2015).

4 Fromme, P., Jordan, P. & Krauß, N. Structure of photosystem I. Biochim. Biophys. Acta, Bioenerg. 1507, 5–31 (2001).

5 Suga, M. & Shen, J.-R. Structural variations of photosystem I-antenna supercomplex in response to adaptations to different light environments. Curr. Opin. Struct. Biol. 63, 10–17 (2020).

6 Hippler, M. & Nelson, N. The plasticity of photosystem I. Plant Cell Physiol. 62, 1073–1081 (2021).

7 Jordan, P. et al. Three-dimensional structure of cyanobacterial photosystem I at 2.5 Å resolution. Nature 411, 909–917 (2001).

8 Malavath, T., Caspy, I., Netzer-El, S. Y., Klaiman, D. & Nelson, N. Structure and function of wild-type and subunit-depleted photosystem I in *Synechocystis*. Biochim. Biophys. Acta, Bioenerg. 1859, 645–654 (2018).

9 Dobson, Z. et al. The structure of photosystem I from a high-light-tolerant cyanobacteria. eLife 10, e67518 (2021).

10 Nagao, R. Handbook of cyanobacterial PSI structures. ([Kindle edition]. Retrieved from Amazon.com, 2022).

11 Kato, K. et al. Structure of a cyanobacterial photosystem I tetramer revealed by cryo-electron microscopy. Nat. Commun. 10, 4929 (2019).

12 Zheng, L. et al. Structural and functional insights into the tetrameric photosystem I from heterocyst-forming cyanobacteria. Nat. Plants 5, 1087–1097 (2019).

13 Chen, M. et al. Distinct structural modulation of photosystem I and lipid environment stabilizes its tetrameric assembly. Nat. Plants 6, 314–320 (2020).

14 Semchonok, D. A. et al. Cryo-EM structure of a tetrameric photosystem I from *Chroococcidiopsis* TS-821, a thermophilic, unicellular, non-heterocyst-forming cyanobacterium. Plant Commun. 3, 100248 (2022).

15 Watanabe, M., Kubota, H., Wada, H., Narikawa, R. & Ikeuchi, M. Novel supercomplex organization of photosystem I in *Anabaena* and *Cyanophora paradoxa*. Plant Cell Physiol. 52, 162–168 (2011).

16 Li, M. et al. Physiological and evolutionary implications of tetrameric photosystem I in cyanobacteria. Nat. Plants 5, 1309–1319 (2019).

17 Falkowski, P. G. et al. The evolution of modern eukaryotic phytoplankton. Science 305, 354–360 (2004).

18 Laudenbach, D. E. & Straus, N. A. Characterization of a cyanobacterial iron stress-induced gene similar to *psbC*. J. Bacteriol. 170, 5018–5026 (1988).

19 Burnap, R. L., Troyan, T. & Sherman, L. A. The highly abundant chlorophyll-protein complex of iron-deficient *Synechococcus* sp. PCC7942 (CP43’) is encoded by the *isiA* gene. Plant Physiol. 103, 893–902 (1993).

20 Jia, A., Zheng, Y., Chen, H. & Wang, Q. Regulation and functional complexity of the chlorophyll-binding protein IsiA. Front. Microbiol. 12, 774107 (2021).

21 Bibby, T. S., Nield, J. & Barber, J. Iron deficiency induces the formation of an antenna ring around trimeric photosystem I in cyanobacteria. Nature 412, 743–745 (2001).

22 Boekema, E. J. et al. A giant chlorophyll-protein complex induced by iron deficiency in cyanobacteria. Nature 412, 745–748 (2001).

23 Toporik, H., Li, J., Williams, D., Chiu, P.-L. & Mazor, Y. The structure of the stress-induced photosystem I-IsiA antenna supercomplex. Nat. Struct. Mol. Biol. 26, 443–449 (2019).

24 Cao, P. et al. Structural basis for energy and electron transfer of the photosystem I-IsiA-flavodoxin supercomplex. Nat. Plants 6, 167–176 (2020).

25 Akita, F. et al. Structure of a cyanobacterial photosystem I surrounded by octadecameric IsiA antenna proteins. Commun. Biol. 3, 232 (2020).

26 Andrizhiyevskaya, E. G. et al. Spectroscopic properties of PSI-IsiA supercomplexes from the cyanobacterium *Synechococcus* PCC 7942. Biochim. Biophys. Acta, Bioenerg. 1556, 265–272 (2002).

27 Melkozernov, A. N., Bibby, T. S., Lin, S., Barber, J. & Blankenship, R. E. Time-resolved absorption and emission show that the CP43’ antenna ring of iron-stressed *Synechocystis* sp. PCC6803 is efficiently coupled to the photosystem I reaction center core. Biochemistry 42, 3893–3903 (2003).

28 Chen, H.-Y. S., Liberton, M., Pakrasi, H. B. & Niedzwiedzki, D. M. Reevaluating the mechanism of excitation energy regulation in iron-starved cyanobacteria. Biochim. Biophys. Acta, Bioenerg. 1858, 249–258 (2017).

29 Shen, G., Gan, F. & Bryant, D. A. The siderophilic cyanobacterium *Leptolyngbya* sp. strain JSC-1 acclimates to iron starvation by expressing multiple isiA-family genes. Photosynth. Res. 128, 325–340 (2016).

30 Nagao, R. et al. Molecular organizations and function of iron-stress-induced-A protein family in *Anabaena* sp. PCC 7120. Biochim. Biophys. Acta, Bioenerg. 1862, 148327 (2021).

31 Nagao, R. et al. Excitation-energy transfer in heterocysts isolated from the cyanobacterium *Anabaena* sp. PCC 7120 as studied by time-resolved fluorescence spectroscopy. Biochim. Biophys. Acta, Bioenerg. 1863, 148509 (2021).

32 Nagao, R. et al. pH-induced regulation of excitation energy transfer in the cyanobacterial photosystem I tetramer. J. Phys. Chem. B 124, 1949–1954 (2020).

33 Chitnis, V. P. & Chitnis, P. R. PsaL subunit is required for the formation of photosystem I trimers in the cyanobacterium *Synechocystis* sp. PCC 6803. FEBS Lett. 336, 330–334 (1993).

34 Schluchter, W. M., Shen, G., Zhao, J. & Bryant, D. A. Characterization of *psaI* and *psaL* mutants of *Synechococcus* sp. strain PCC 7002: a new model for state transitions in cyanobacteria. Photochem. Photobiol. 64, 53–66 (1996).

35 Li, M., Semchonok, D. A., Boekema, E. J. & Bruce, B. D. Characterization and evolution of tetrameric photosystem I from the thermophilic cyanobacterium *Chroococcidiopsis* sp TS-821. Plant Cell 26, 1230–1245 (2014).

36 Park, Y.-I., Sandström, S., Gustafsson, P. & Öquist, G. Expression of the *isiA* gene is essential for the survival of the cyanobacterium *Synechococcus* sp. PCC 7942 by protecting photosystem II from excess light under iron limitation. Mol. Microbiol. 32, 123–129 (1999).

37 Wang, Q., Hall, C. L., Al-Adami, M. Z. & He, Q. IsiA Is required for the formation of photosystem I supercomplexes and for efficient state transition in *Synechocystis* PCC 6803. PLoS One 5 (2010).

38 Kato, K. et al. Structural basis for the absence of low-energy chlorophylls in a photosystem I trimer from *Gloeobacter violaceus*. eLife 11, e73990 (2022).

39 Nagao, R., Yamaguchi, M., Nakamura, S., Ueoka-Nakanishi, H. & Noguchi, T. Genetically introduced hydrogen bond interactions reveal an asymmetric charge distribution on the radical cation of the special-pair chlorophyll P680. J. Biol. Chem. 292, 7474–7486 (2017).

40 Polívka, T. & Sundström, V. Ultrafast dynamics of carotenoid excited states-From solution to natural and artificial systems. Chem. Rev. 104, 2021–2072 (2004).

41 Ruban, A. V., Johnson, M. P. & Duffy, C. D. P. The photoprotective molecular switch in the photosystem II antenna. Biochim. Biophys. Acta, Bioenerg. 1817, 167–181 (2012).

42 Pinto, F. L., Thapper, A., Sontheim, W. & Lindblad, P. Analysis of current and alternative phenol based RNA extraction methodologies for cyanobacteria. BMC Mol. Biol. 10, 79 (2009).

43 Ehira, S. & Miyazaki, S. Regulation of genes involved in heterocyst differentiation in the cyanobacterium *Anabaena* sp. strain PCC 7120 by a group 2 sigma factor SigC. Life 5, 587–603 (2015).

44 Ikeuchi, M. & Inoue, Y. A new photosystem II reaction center component (4.8 kDa protein) encoded by chloroplast genome. FEBS Lett. 241, 99–104 (1988).

45 Nagao, R. et al. Structural basis for energy harvesting and dissipation in a diatom PSII-FCPII supercomplex. Nat. Plants 5, 890–901 (2019).

46 Schägger, H. & von Jagow, G. Blue native electrophoresis for isolation of membrane protein complexes in enzymatically active form. Anal. Biochem. 199, 223–231 (1991).

47 Nagao, R., Yokono, M., Akimoto, S. & Tomo, T. High excitation energy quenching in fucoxanthin chlorophyll *a*/*c*-binding protein complexes from the diatom *Chaetoceros gracilis*. J. Phys. Chem. B 117, 6888–6895 (2013).

48 Hamada, F., Murakami, A. & Akimoto, S. Adaptation of divinyl chlorophyll *a*/*b*-containing cyanobacterium to different light conditions: three strains of *Prochlorococcus marinus*. J. Phys. Chem. B 121, 9081–9090 (2017).

49 Nagao, R., Yokono, M., Ueno, Y., Shen, J.-R. & Akimoto, S. Excitation-energy transfer and quenching in diatom PSI-FCPI upon P700 cation formation. J. Phys. Chem. B 124, 1481–1486 (2020).

50 Yonekura, K., Maki-Yonekura, S., Naitow, H., Hamaguchi, T. & Takaba, K. Machine learning-based real-time object locator/evaluator for cryo-EM data collection. Commun. Biol. 4, 1044 (2021).

51 Mastronarde, D. N. Automated electron microscope tomography using robust prediction of specimen movements. J. Struct. Biol. 152, 36–51 (2005).

52 Zheng, S. Q. et al. MotionCor2: anisotropic correction of beam-induced motion for improved cryo-electron microscopy. Nat. Methods 14, 331–332 (2017).

53 Mindell, J. A. & Grigorieff, N. Accurate determination of local defocus and specimen tilt in electron microscopy. J. Struct. Biol. 142, 334–347 (2003).

54 Zivanov, J., Nakane, T. & Scheres, S. H. W. Estimation of high-order aberrations and anisotropic magnification from cryo-EM data sets in *RELION-3.1*. IUCrJ 7, 253–267 (2020).

55 Grigorieff, N. & Harrison, S. C. Near-atomic resolution reconstructions of icosahedral viruses from electron cryo-microscopy. Curr. Opin. Struc. Biol. 21, 265–273 (2011).

56 Bepler, T., Kelley, K., Noble, A. J. & Berger, B. Topaz-Denoise: general deep denoising models for cryoEM and cryoET. Nat. Commun. 11, 5208 (2020).

57 Kelley, L. A., Mezulis, S., Yates, C. M., Wass, M. N. & Sternberg, M. J. E. The Phyre2 web portal for protein modeling, prediction and analysis. Nat. Protoc. 10, 845–858 (2015).

58 Pettersen, E. F. et al. UCSF chimera - A visualization system for exploratory research and analysis. J. Comput. Chem. 25, 1605–1612 (2004).

59 Emsley, P., Lohkamp, B., Scott, W. G. & Cowtan, K. Features and development of *Coot*. Acta Crystallogr. D Biol. Crystallogr. 66, 486–501 (2010).

60 Adams, P. D. et al. PHENIX: a comprehensive Python-based system for macromolecular structure solution. Acta Crystallogr. D Biol. Crystallogr. 66, 213–221 (2010).

61 Yamashita, K., Palmer, C. M., Burnley, T. & Murshudov, G. N. Cryo-EM single-particle structure refinement and map calculation using Servalcat. Acta Crystallogr. D Struct. Biol. 77, 1282–1291 (2021).

62 Chen, V. B. et al. MolProbity: all-atom structure validation for macromolecular crystallography. Acta Crystallogr. D Biol. Crystallogr. 66, 12–21 (2010).

63 Barad, B. A. et al. EMRinger: side chain-directed model and map validation for 3D cryo-electron microscopy. Nat. Methods 12, 943–946 (2015).

64 Pintilie, G. et al. Measurement of atom resolvability in cryo-EM maps with *Q*-scores. Nat. Methods 17, 328–334 (2020).

65 Schrödinger, L. L. C. The PyMOL Molecular Graphics System. Version 2.5.0. (2021).

