## Supplementary Information for "Structure of a monomeric photosystem I core associated with iron-stress-induced-A proteins from *Anabaena* sp. PCC 7120"

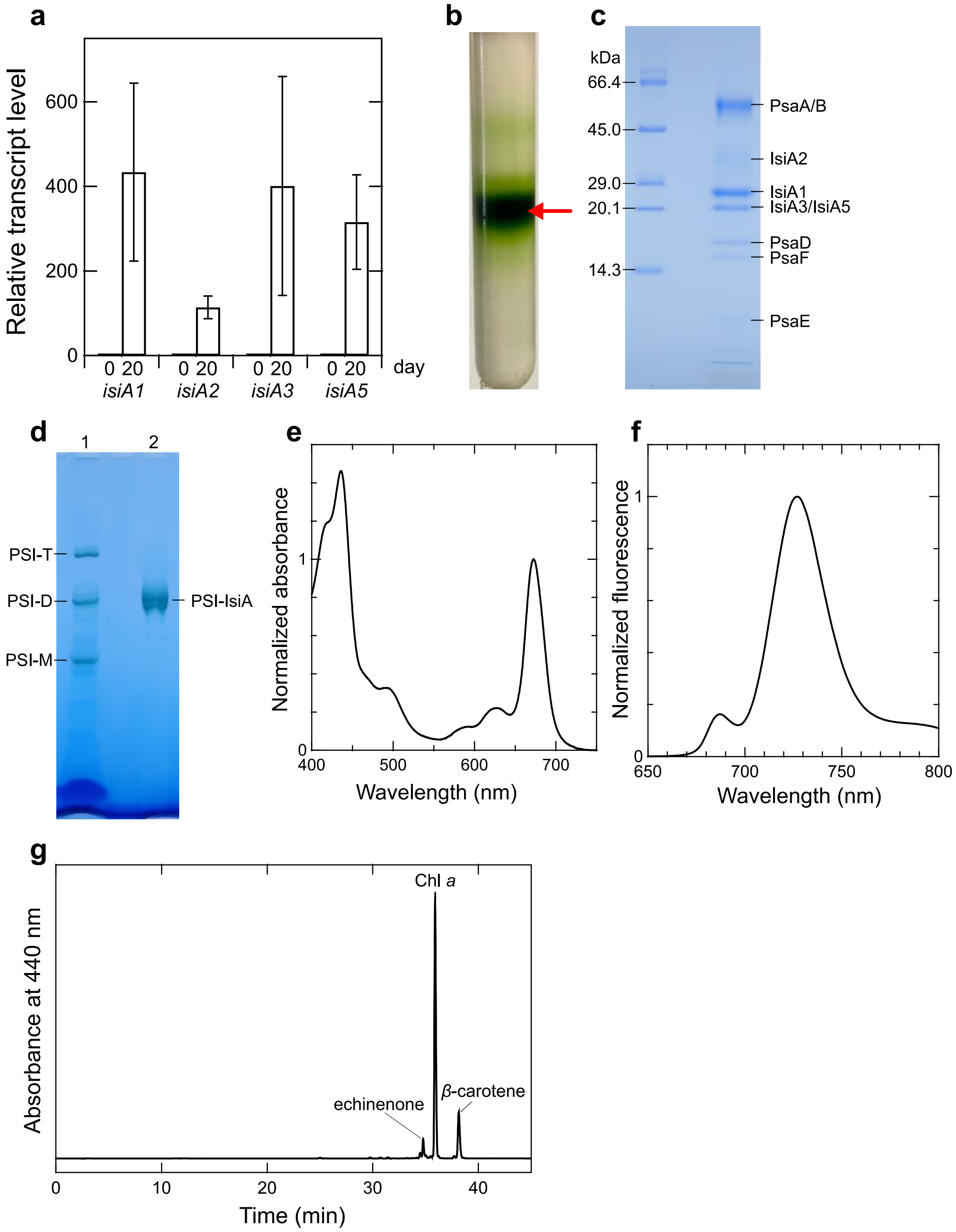


**Supplementary Fig. 1. Expression of the *isiA* genes, and purification, characterization of the *Anabaena* PSI-IsiA supercomlex.**

**a,** Expression of the *isiA* genes. The transcript levels of *isiA1*, *isiA2*, *isiA3*, and *isiA5* were determined by qRT-PCR for cells grown under the iron-deficient condition for 0 or 20 days. The transcript level for each gene before incubation of the iron deficiency (0 day) was taken as 1. RNA samples were prepared from four independently grown cultures. Bars and error bars represent means ± S.D. (n = 4). **b,** Trehalose density gradient centrifugation. The red arrow indicates the PSI-IsiA fraction. **c,** SDS-PAGE analysis of PSI-IsiA. Each band was assigned by mass spectrometry analysis. **d,** BN-PAGE analysis of PSI-IsiA. Lane 1, thylakoid membranes as a reference; lane 2, PSI-IsiA. PSI-T, PSI-D, and PSI-M correspond to PSI tetramer, PSI dimer, and PSI monomer, respectively. **e,** Absorption spectrum of PSI-IsiA measured at 298 K. Three measurements were averaged, and the resultant spectrum was normalized by the intensity of the Qy peak. **f,** Fluorescence-emission spectrum of PSI-IsiA measured at 77 K upon excitation at 430 nm. Three measurements were averaged, and the resultant spectrum was normalized by the intensity of the PSI fluorescence at 727 nm. **g,** HPLC analysis of the pigments extracted from PSI-IsiA monitored at 440 nm. Data in panels **b**, **c**, **d,** and **g** are representative of three independent experiments.

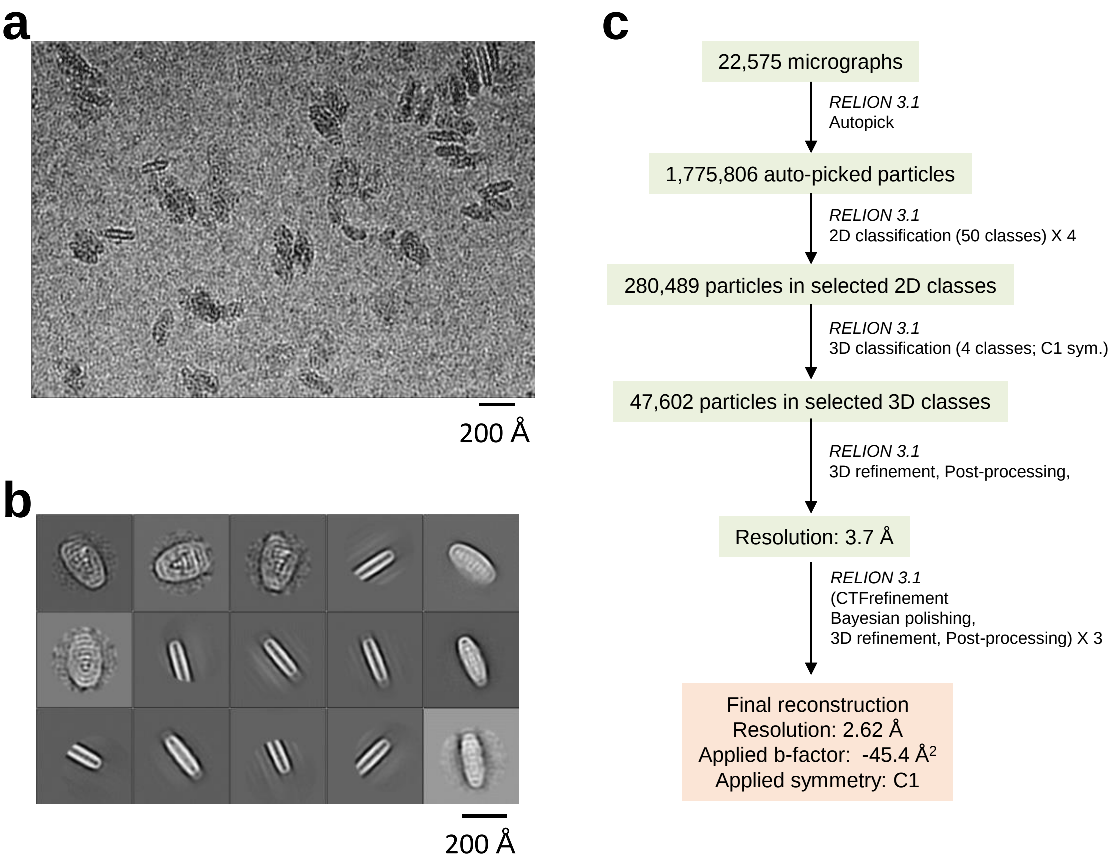


**Supplementary Fig. 2. Cryo-EM data collection and processing of PSI-IsiA.**

**a,** A representative cryo-EM micrograph of PSI-IsiA. **b,** Representative 2D classes of PSI-IsiA. The box size is 396 Å. **c,** Schematic flowchart showing the classification scheme and data processing for PSI-IsiA. The overall PSI-IsiA structure was reconstructed at 2.62-Å resolution from 47,602 particles. See Methods section for more details.

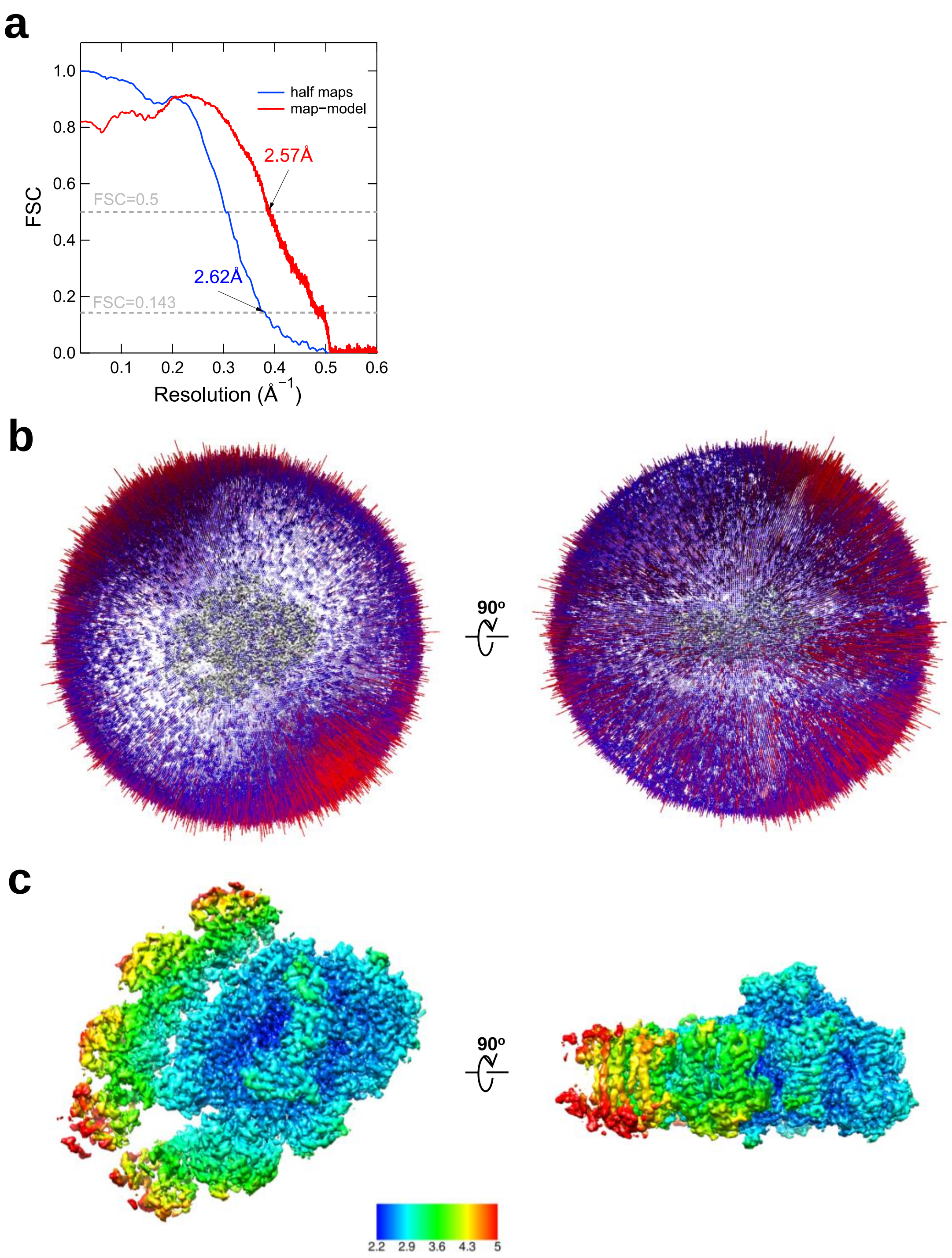


**Supplementary Fig. 3. Evaluation of the cryo-EM map quality.**

**a,** FSC curves of PSI-IsiA for independently refined half maps (blue) and for map-minus-model (red). **b,** Angular distributions of the particles used for the reconstruction of PSI-IsiA. Each cylinder represents one view, and the height of the cylinder is proportional to the number of particles for that view. **c,** Local resolution maps of PSI-IsiA.

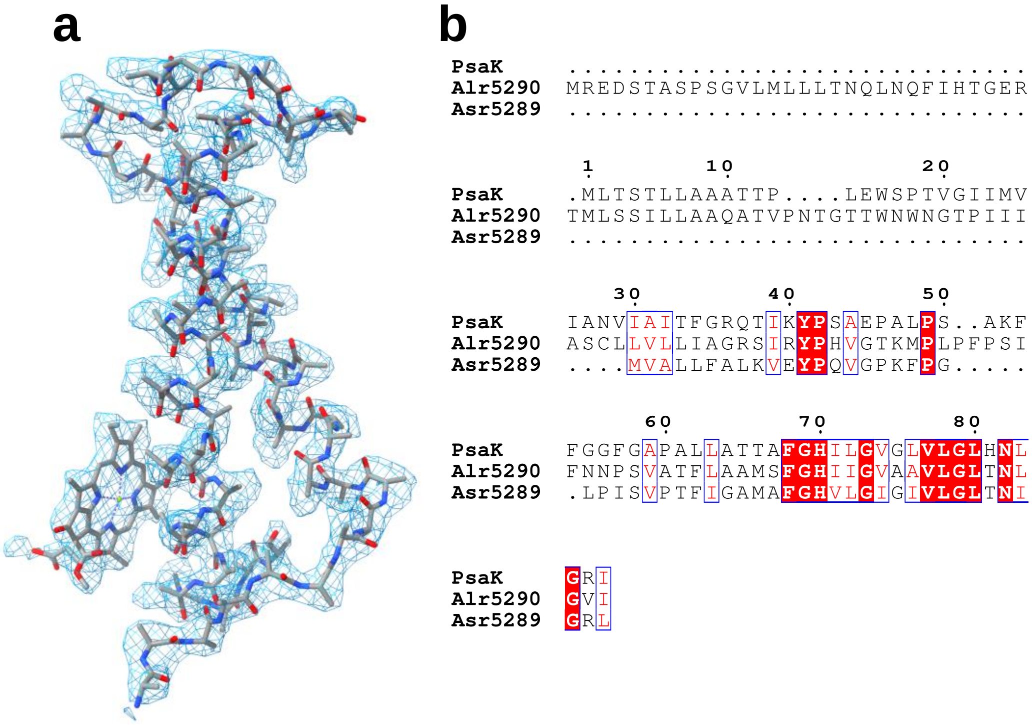


**Supplementary Fig. 4. Evaluation of the position for PsaK.**

**a,** The density for Unknown and its corresponding model are shown as blue meshes and gray sticks, respectively. **b,** Multiple sequence alignment (ClustalW and ESPript) among three PsaK family proteins (PsaK, Alr5290, Asr5289) in *Anabaena*.

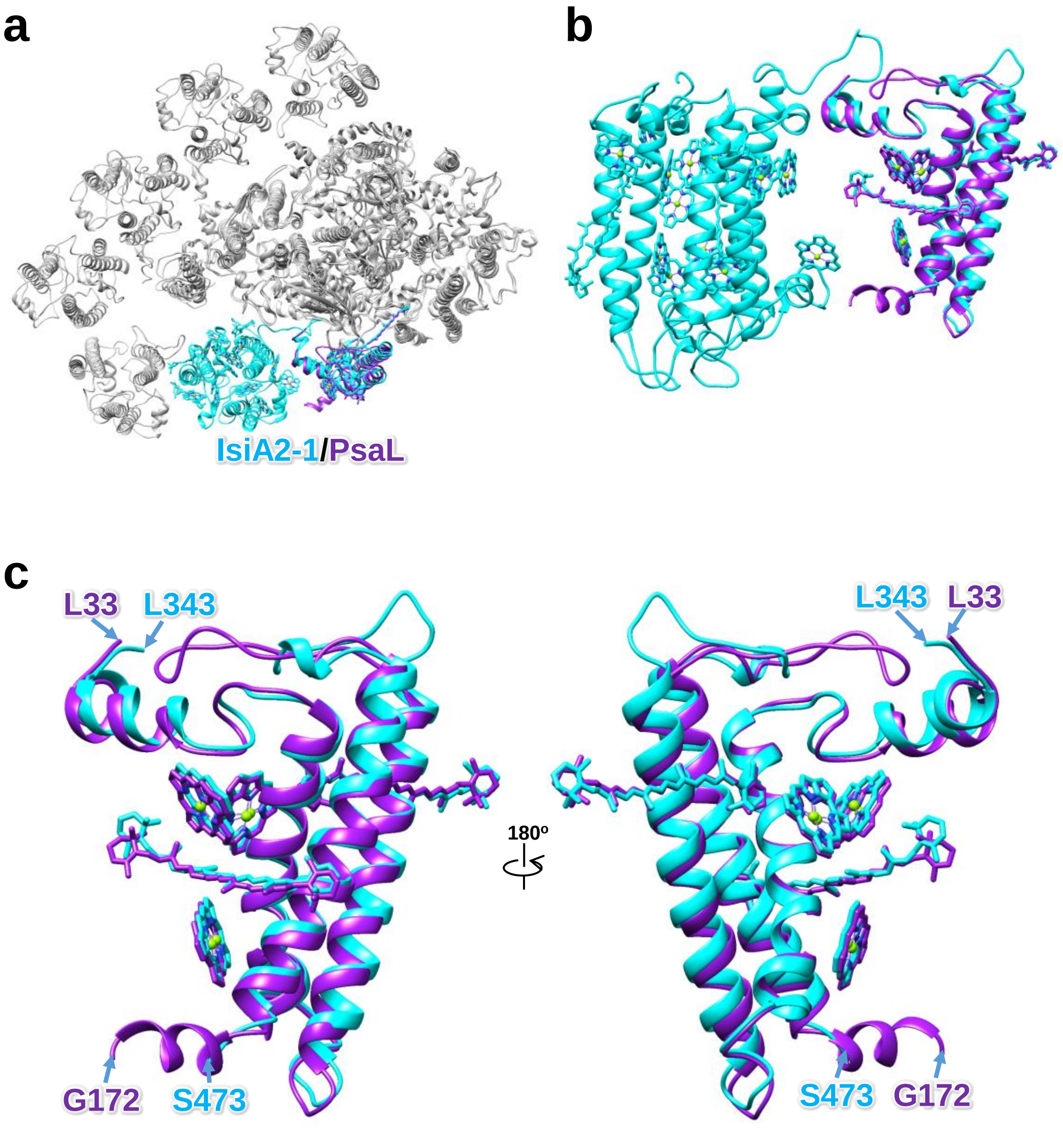
a
b
c
IsiA2-1/PsaL
L33
L343
G172
S473
180º
L343
L33
S473
G172

**d**

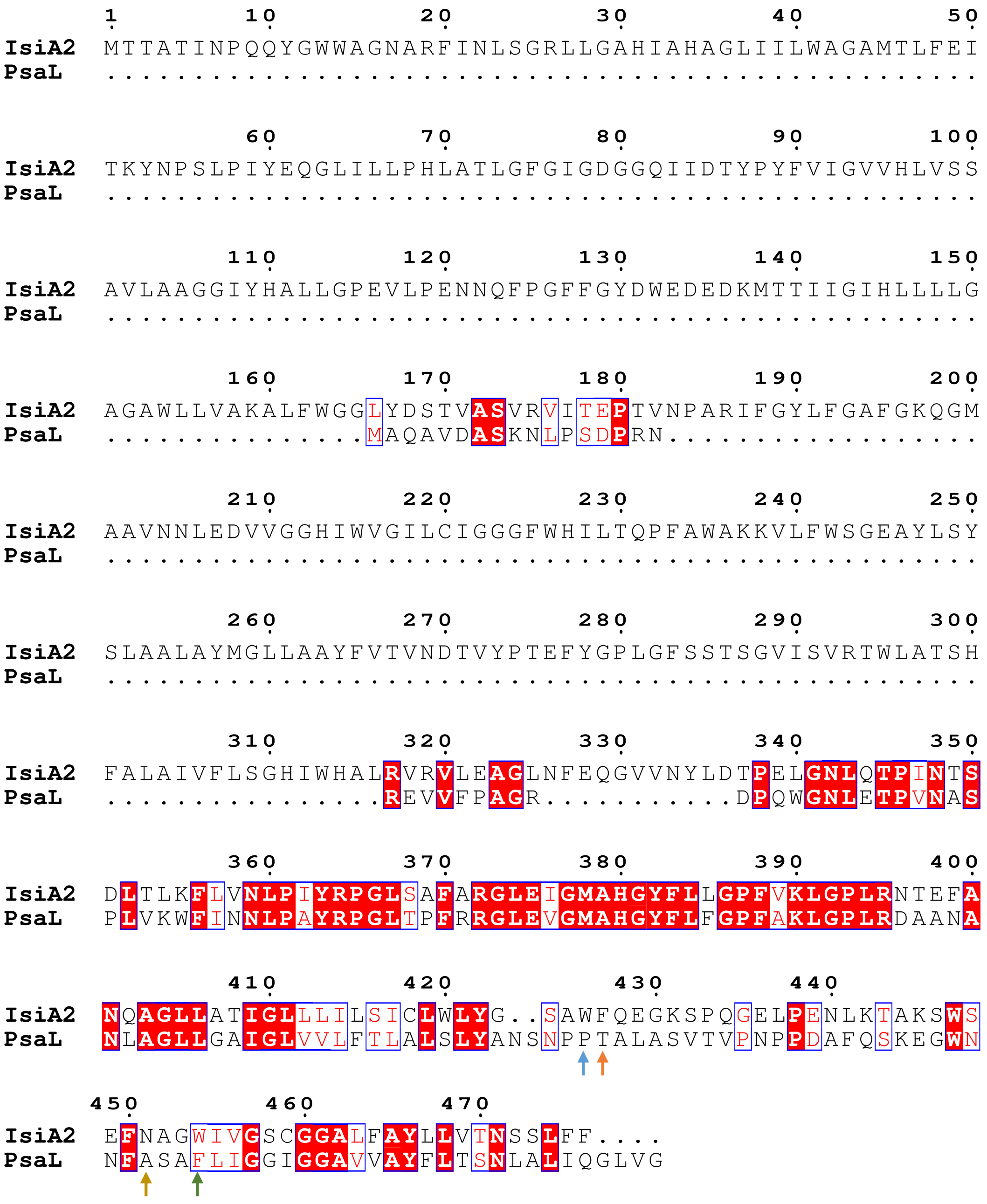
IsiA2 MTTATINPQQYGWWAGNARFINLSGRLLGAHIAHAGLIILWAGAMTLFEI
PsaL ..................................................
IsiA2 TKYNPSLPIYEQGLILLPHLATLGFGIGDGGQIIDTYPYFVIGVVHLVSS
PsaL ..................................................
IsiA2 AVLAAGGIYHALLGPEVLPENNQFPGFFGYDWEDEDKMTTIIGIHLLLLG
PsaL ..................................................
IsiA2 AGAWLLVAKALFWGGLYDSTVASVRVITEPTVNPARIFGYLFGAFGKQGM
PsaL ...............MAQAVDASKNLPSDPRN..................
IsiA2 AAVNNLEDVVGGHIWVGILCIGGGFWHILTQPFAWAKKVLFWSGEAYLSY
PsaL ..................................................
IsiA2 SLAALAYMGLLAAYFVTVNDTVYPTEFYGPLGFSSTSGVISVRTWLATSH
PsaL ..................................................
IsiA2 FALAIVFLSGHIWHALRVRVLEAGLNFEQGVVNYLDTPELGNLQTPINTS
PsaL ................REVVFPAGR...........DPQWGNLETPVNAS
IsiA2 DLTLKFLVNLPIYRPGLSAFARGLEIGMAHGYFLLGPFVKLGPLRNTEFA
PsaL PLVKWFINNLPAYRPGLTPFRRGLEVGMAHGYFLFGPFAKLGPLRDAANA
IsiA2 NQAGLLATIGLLLILSICLWLYG..SAWFQEGKSPQGELPENLKTAKSWS
PsaL NLAGLLGAIGLVVLFTLALSLYANSNPPTALASVTVPNPPDAFQSKEGWN
IsiA2 EFNAGWIVGSCGGALFAYLLVTNSSLFF....
PsaL NFASAFLIGGIGGAVVAYFLTSNLALIQGLVG

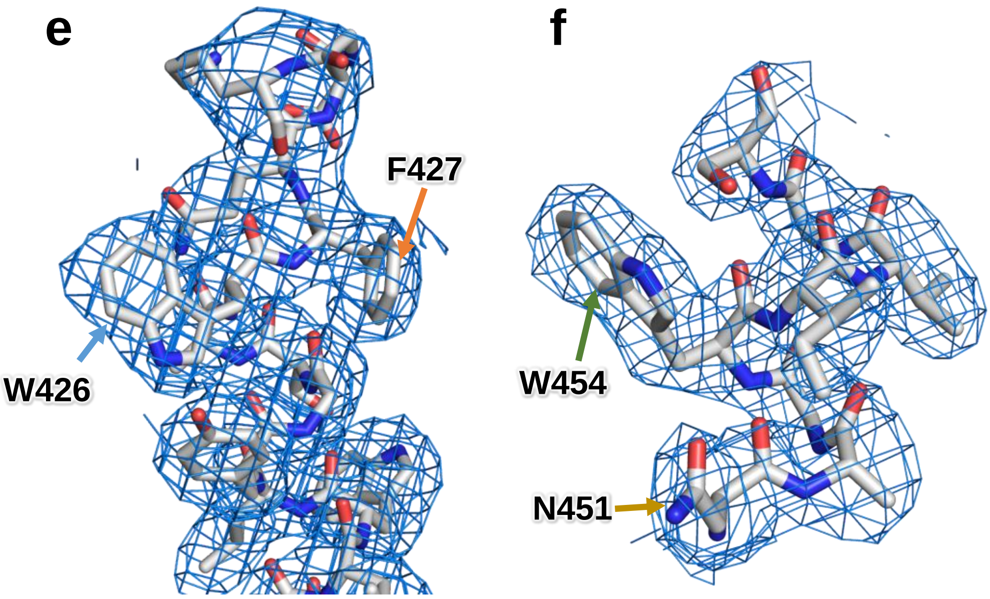


**Supplementary Fig. 5. Comparison between IsiA2-1 and PsaL.**

**a,** Superposition of the PSI-IsiA structure with the PSI-monomer structure prepared from the *Anabaena* PSI tetramer (PDB: 6JEO). The structures are viewed from the cytosolic side. IsiA2-1 and PsaL are colored cyan and purple, respectively, whereas the other subunits are colored gray. **b,** Side view of the superposition of structures between IsiA2-1 and PsaL. **c,** Structural comparison of Cα atoms, Chls, and *β*-carotenes between the C-terminal domain of IsiA2 and PsaL. Only rings of the Chl molecules are depicted in panels **a**–**c**. **d,** Multiple sequence alignment (ClustalW and ESPript) between IsiA2 and PsaL in *Anabaena*. Unique residues between IsiA2 and PsaL are indicated by arrows with different colors, which were used for the identification of IsiA2. **e, f,** Characteristic maps and residues of the C-terminal PsaL-like domain of IsiA2. The maps are shown as meshes at 1 σ contour level, and the corresponding models are shown as sticks. The characteristic amino acids shown in panel **c** are labeled with the corresponding arrows.

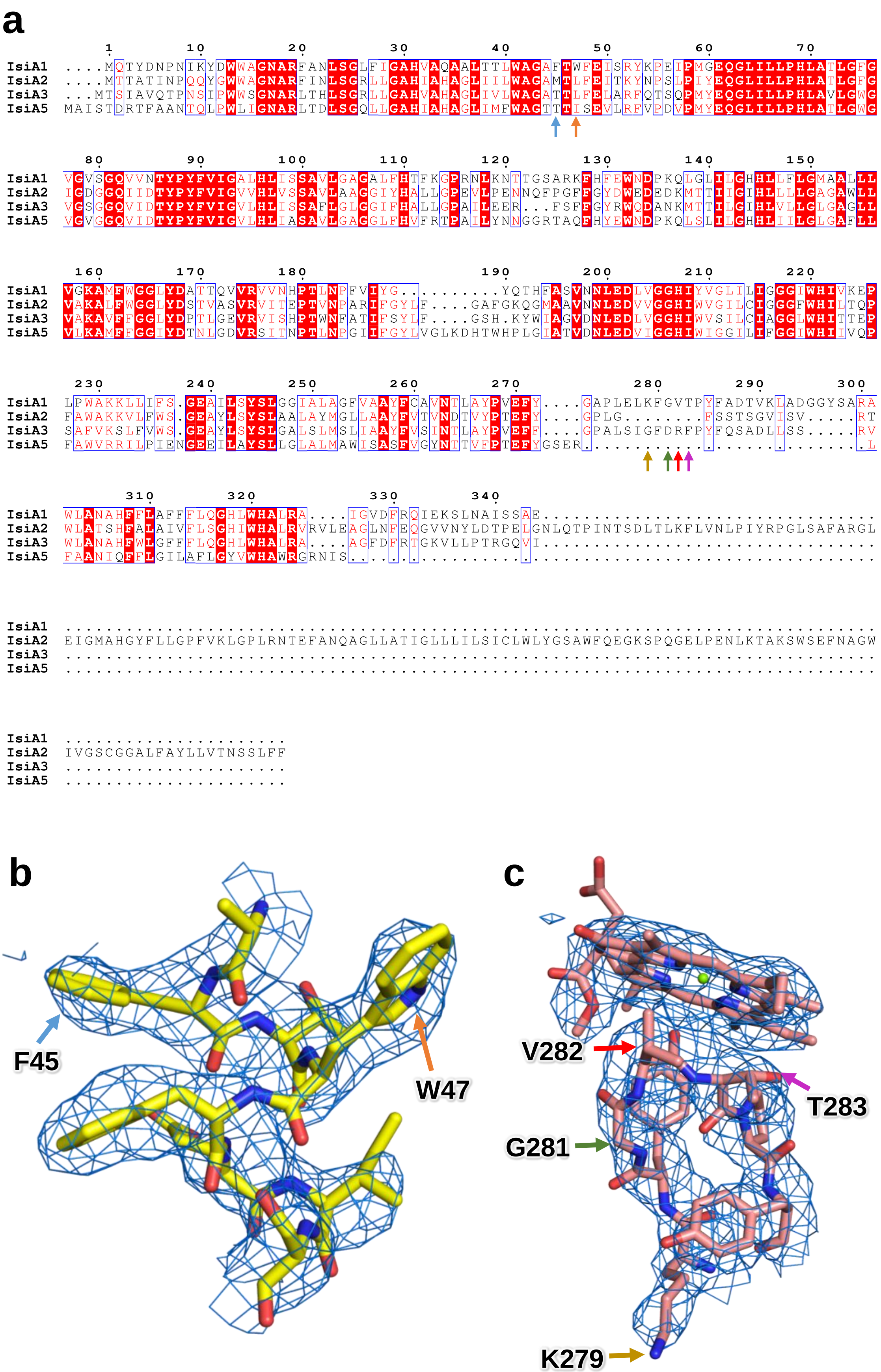
a
IsiA1 ....MQTYDNPNIKYDWWAGNARFANLSGLFIGAHVAQAALTTLWAGAFTWFEISRYKPEIPMGEQGLILLPHLATLGFG
IsiA2 ....MTTATINPQQYGWWAGNARFINLSGRLLGAHIAHAGLIILWAGAMTLFEITKYNPSLPIYEQGLILLPHLATLGFG
IsiA3 ...MTSIAVQTPNSIPWWSGNARLTHLSGRLLGAHVAHAGLIVLWAGATTLFELARFQTSQPMYEQGLILLPHLAVLGWG
IsiA5 MAISTDRTFAANTQLPWLIGNARLTDLSGQLLGAHIAHAGLIMFWAGTTTISEVLRFVPDVPMYEQGLILLPHLATLGWG
IsiA1 VGVSGQVVNTYPYFVIGALHLISSAVLGAGALFHTFKGPRNLKNTTGSARKFHFEWNDPKQLGLILGHHLLFLGMAALLL
IsiA2 IGDGGQIIDTYPYFVIGVVHLVSSAVLAAGGIYHALLGPEVLPENNQFPGFFGYDWEDEDKMTTIIGIHLLLLGAGAWLL
IsiA3 VGSGGQVIDTYPYFVIGVLHLISSAFLGLGGIFHALLGPAILEER...FSFFGYRWQDANKMTTILGIHLVLLGLGAGLL
IsiA5 VGVGGQVIDTYPYFVIGVLHLIASAVLGAGGLFHVFRTPAILYNNGGRTAQFHYEWNDPKQLSLILGHHLIILGLGAFLL
IsiA1 VGKAMFWGGLYDATTQVVRVVNHPTLNPFVIYG.........YQTHFASVNNLEDLVGGHIYVGLILIGGGIWHIVKEP
IsiA2 VAKALFWGGLYDSTVASVRVITEPTVNPARIFGYLF....GAFGKQGMAAVNNLEDVVGGHIWVGILCIGGGFWHILTQP
IsiA3 VAKAVFFGGLYDPTLGEVRVISHPTWNFATIFSYLF....GSH.KYWIAGVDNLEDLVGGHIWVSILCIAGGLWHITTEP
IsiA5 VLKAMFFGGIYDTNLGDVRSITNPTLNPGIIFGYLVGLKDHTWHPLGIATVDNLEDVIGGHIWIGGILIFGGIWHIIVQP
IsiA1 LPWAKKLLIFS.GEAILSYSLGGIALAGFVAAYFCAVNTLAYPVEFY....GAPLELKFGVTPYFADTVKLADGGYSARA
IsiA2 FAWAKKVLFWS.GEAYLSYSLAALAYMGLLAAYFVTVNDTVYPTEFY....GPLG........FSSTSGVISV.....RT
IsiA3 SAFVKSLFVWS.GEAYLSYSLGALSLMSLIAAYFVSINTLAYPVEFF....GPALSIGFDRFPYFQSADLLSS.....RV
IsiA5 FAWVRRILPIENGEEILAYSLLGLALMAWISASFVGYNTTVEPTEFYGSER..........................L
IsiA1 WLANAHFFLAFFFLQGHLWHALRA....IGVDFRQIEKSLNAISSAE.................................
IsiA2 WLATSHFALAIVFLSGHIWHALRVRVLEAGLNFEQGVVNYLDTPELGNLQTPINTSDLTLKFLVNLPIYRPGLSAFARGL
IsiA3 WLANAHFWLGFFFLQGHLWHALRA....AGFDFRTGKVLLPTRGQVI.................................
IsiA5 FAANIQFFLGILAFLGYVWHAWRGRNIS....................................................
IsiA2 EIGMAHGYFLLGPFVKLGPLRNTEFANQAGLLATIGLLLILSICLWLYGSAWFQEGKSPQGELPENLKTAKSWSEFNAGW
IsiA2 IVGSCGGALFAYLLVTNSSLFF
b
F45
W47
c
V282
T283
G281
K279

**Supplementary Fig. 6. Characteristic amino-acid residues used for the identification of IsiA1.**

**a,** Multiple sequence alignment (ClustalW and ESPript) of the four types of IsiAs in *Anabaena*. Unique residues are indicated by arrows with different colors, which were used for the identification of IsiA1. **b, c,** Characteristic maps and residues of IsiA1. The maps are shown as meshes at 1 σ contour level, and the corresponding models are shown as sticks. The characteristic amino acids shown in panel **a** are labeled with the corresponding arrows.

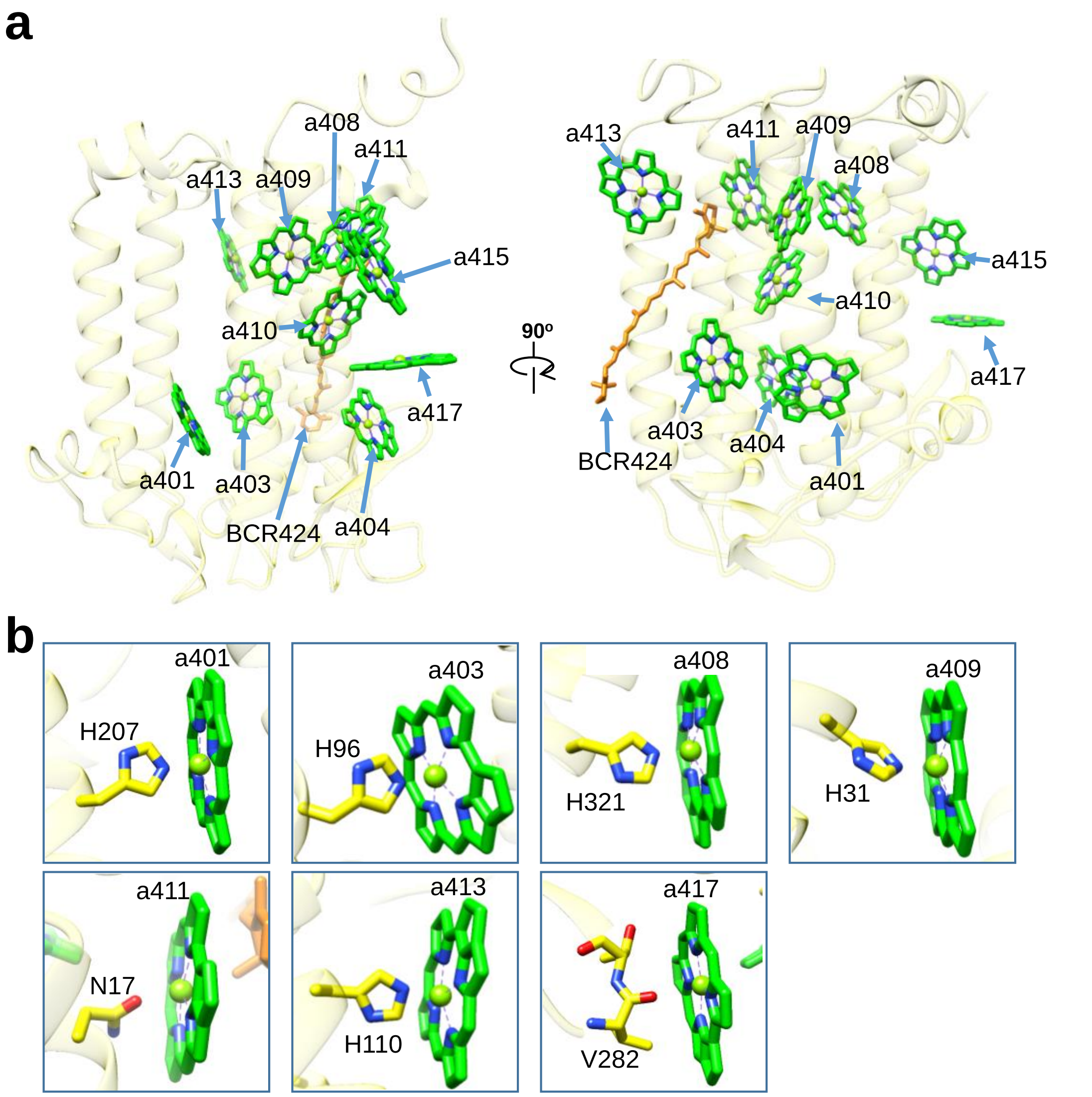


**Supplementary Fig. 7. Structure of IsiA1-4.**

**a,** Structure of IsiA1-4 depicted in Cα atoms and arrangements of Chls and $\beta$-carotene (BCR). **b,** Interactions of Chls with their ligands. Chls and $\beta$-carotene are colored green and orange, respectively. Only rings of the Chl molecules are depicted.

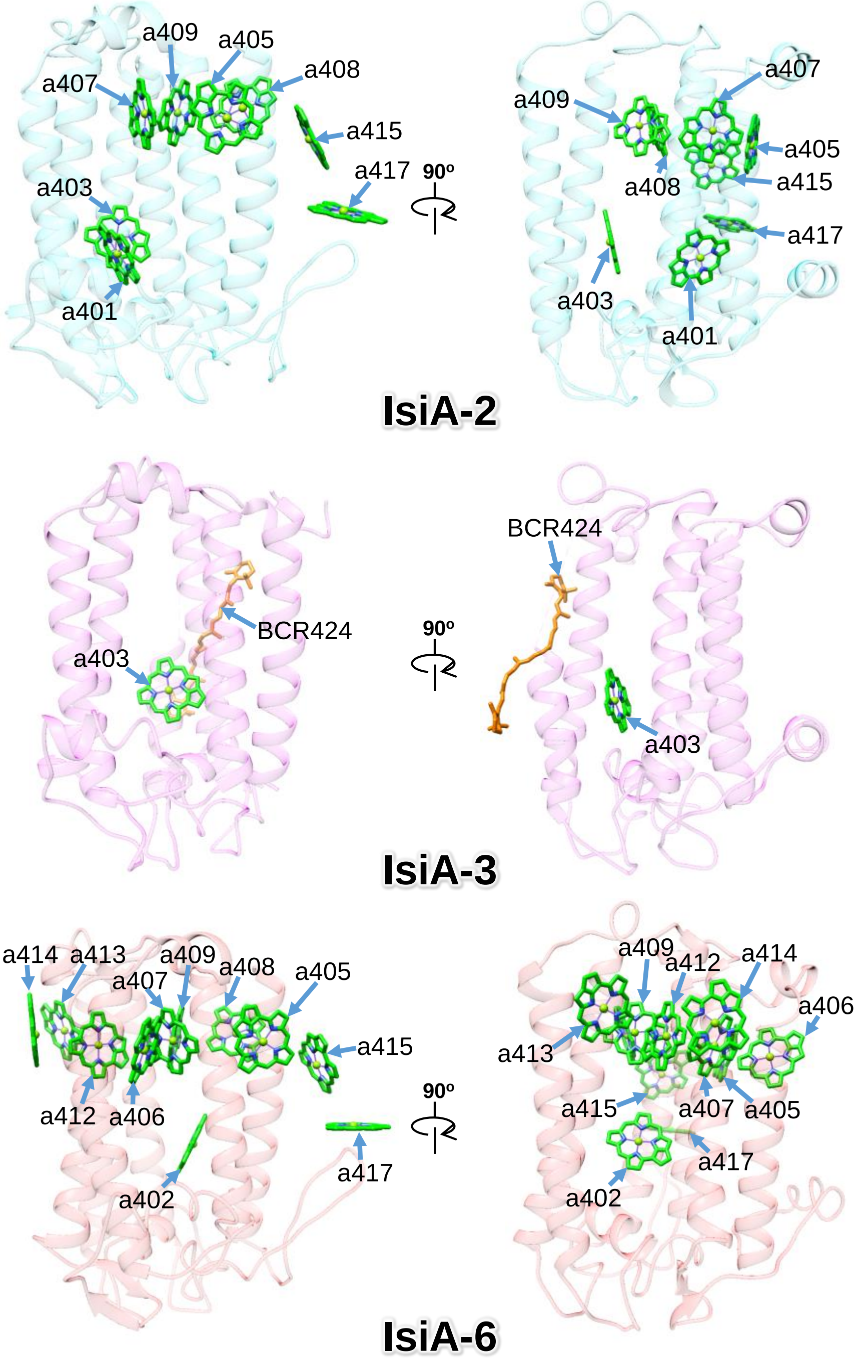


**Supplementary Fig. 8. Structures of IsiA-2, IsiA-3, and IsiA-6.**

Structures of IsiA-2, IsiA-3, and IsiA-6 depicted in Cα atoms and arrangements of Chls and $\beta$-carotene (BCR). Chls and $\beta$-carotene are colored green and orange, respectively. Only rings of the Chl molecules are depicted.

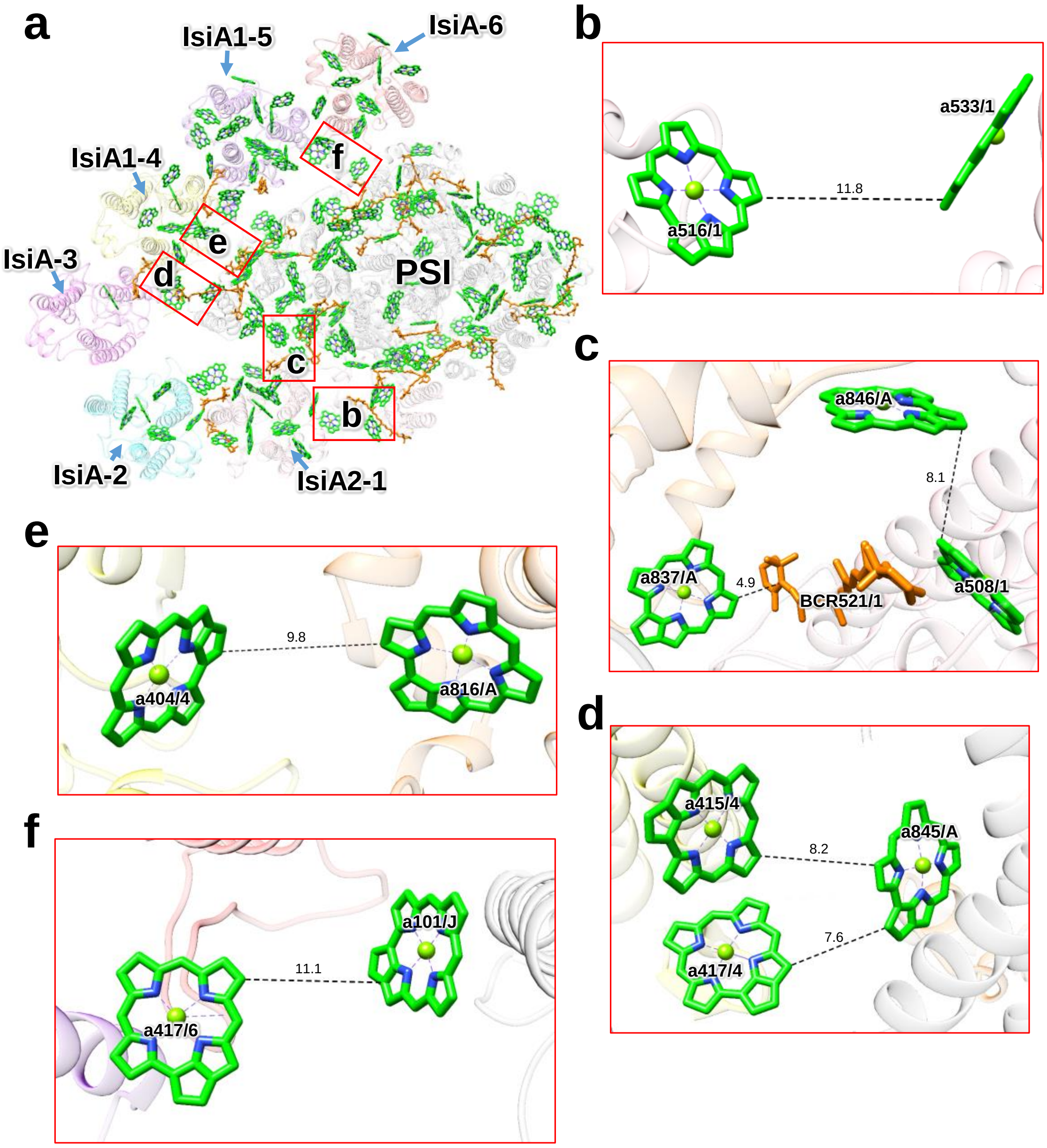


**Supplementary Fig. 9. Pigment-pigment interactions between IsiAs and PSI.**

**a,** The structure of PSI-IsiA viewed from the cytosolic side. Green squared areas are enlarged in panels **b**–**f**. **b**, Interaction between the N-terminal and C-terminal domains within IsiA2-1. **c**, Interactions between IsiA2-1 and PsaA. **d**, **e**, Interactions between IsiA1-4 and PsaA. **f**, Interaction between IsiA-6 and PsaJ. Only rings of the Chl molecules are depicted. Interactions are indicated by dashed lines, and the numbers are distances in Å. The letters represent the numbering of pigments in each subunit of IsiAs/PsaA/PsaJ; for example, a837/A means Chl *a* 837 in PsaA. BCR, $\beta$-carotene.

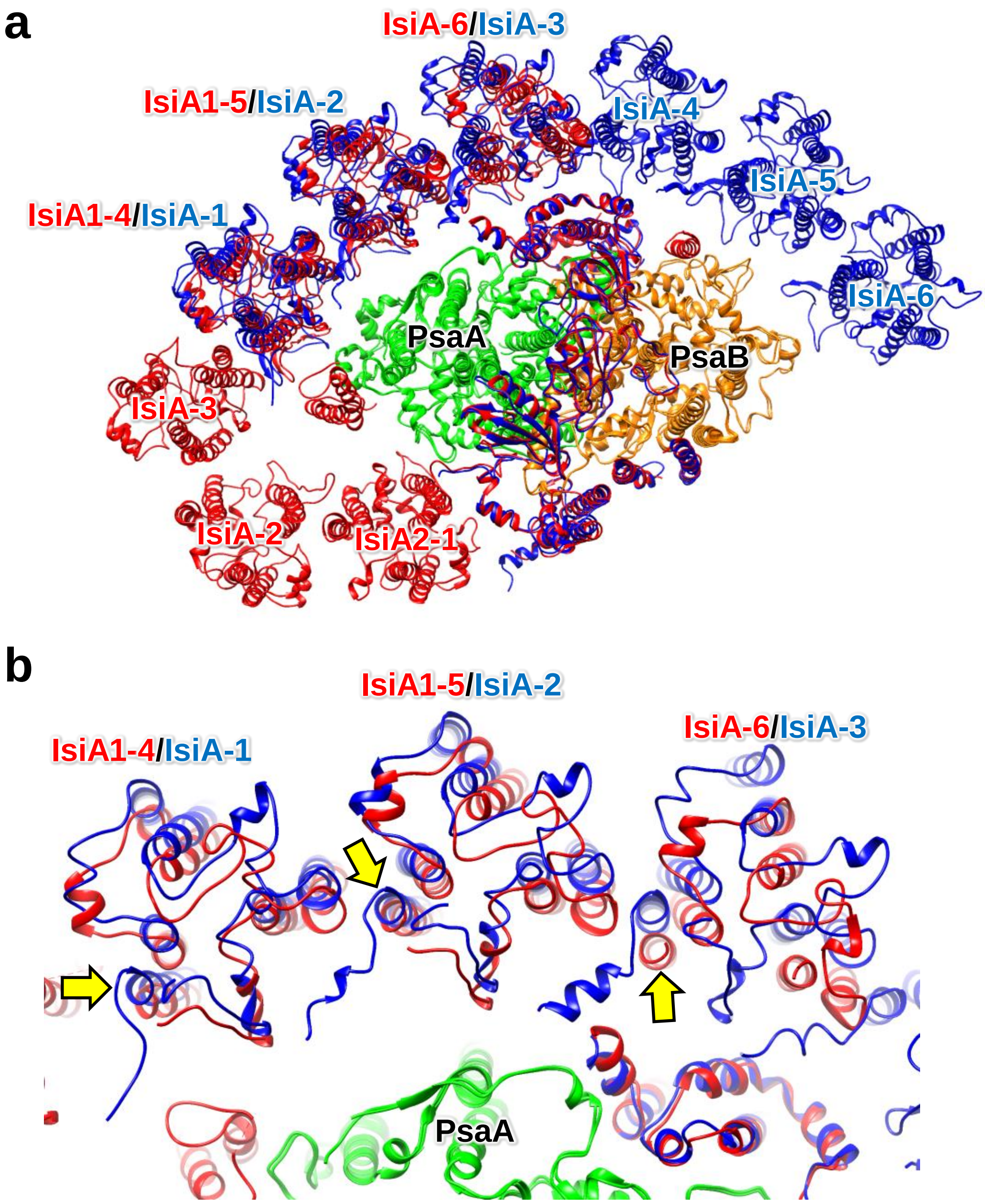
a
IsiA-6/IsiA-3
IsiA1-5/IsiA-2
IsiA-4
IsiA-5
IsiA1-4/IsiA-1
IsiA-6
PsaA
PsaB
IsiA-3
IsiA-2
IsiA2-1
b
IsiA1-5/IsiA-2
IsiA-6/IsiA-3
IsiA1-4/IsiA-1
PsaA

c

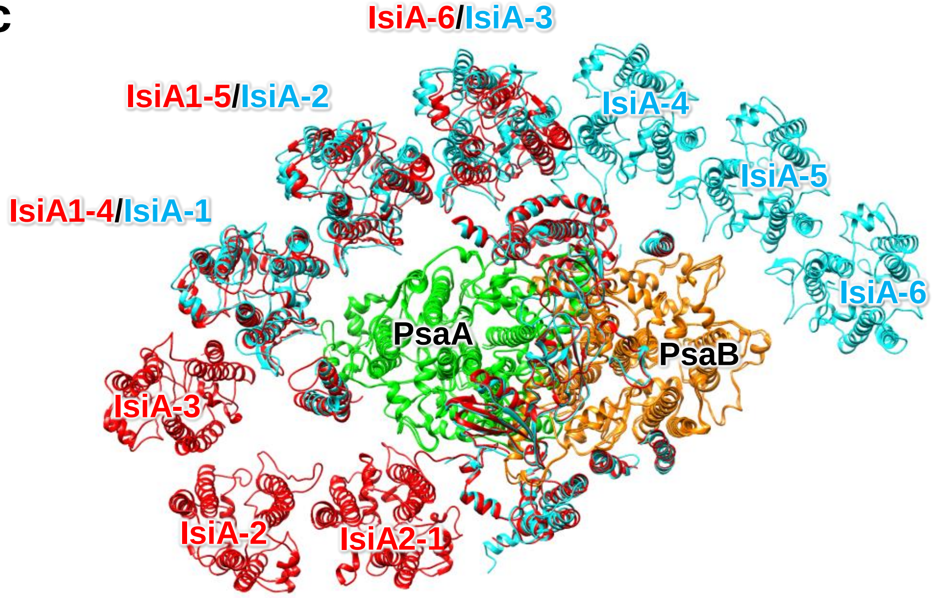


d

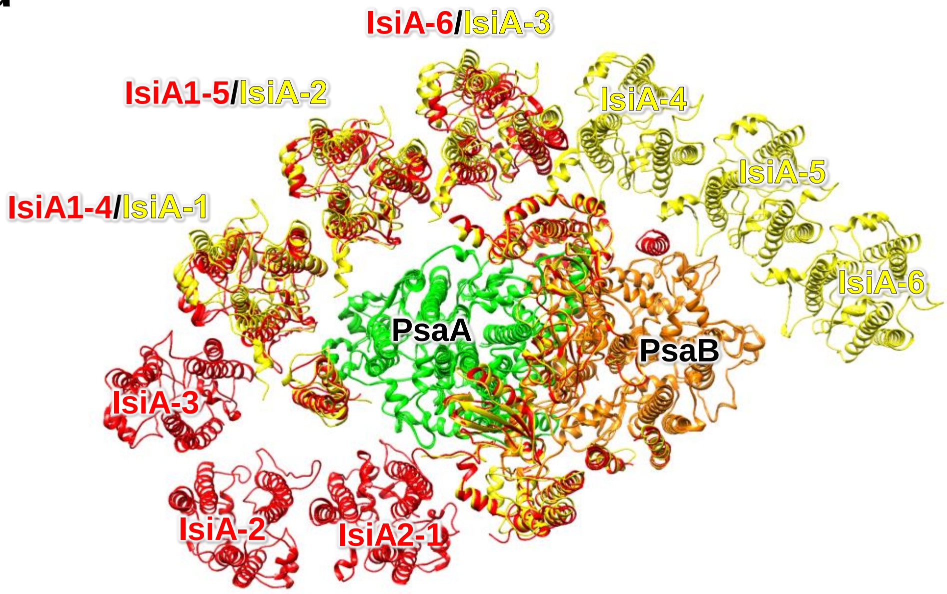


**Supplementary Fig. 10. Structural comparisons of PSI-IsiA between *Anabaena* and other cyanobacteria.**

The structures are viewed from the cytosolic side. The protein structures are PSI-IsiA from *Anabaena*, *Synechocystis* sp. PCC 6803 (**a, b**), *Thermosynechococcus vulcanus* NIES-2134 (**c**), and *Synechococcus elongatus* PCC 7942 (**d**). The subunits of *Anabaena*, *Synechocystis*, *T. vulcanus*, and *Synechococcus* are colored red, blue, cyan, and yellow, respectively, with the colors of PsaA (green) and PsaB (orange) fixed in all the species. The six IsiA subunits in the *Synechocystis*, *T. vulcanus* and *Synechococcus* PSI-IsiA supercomplexes are temporally named as IsiA-1 to IsiA-6. **a, c,** and **d,** overall structures; **b,** expanded view in the interface between characteristic IsiAs and PSI. The C-terminal loops of each IsiA are indicated by yellow arrows.

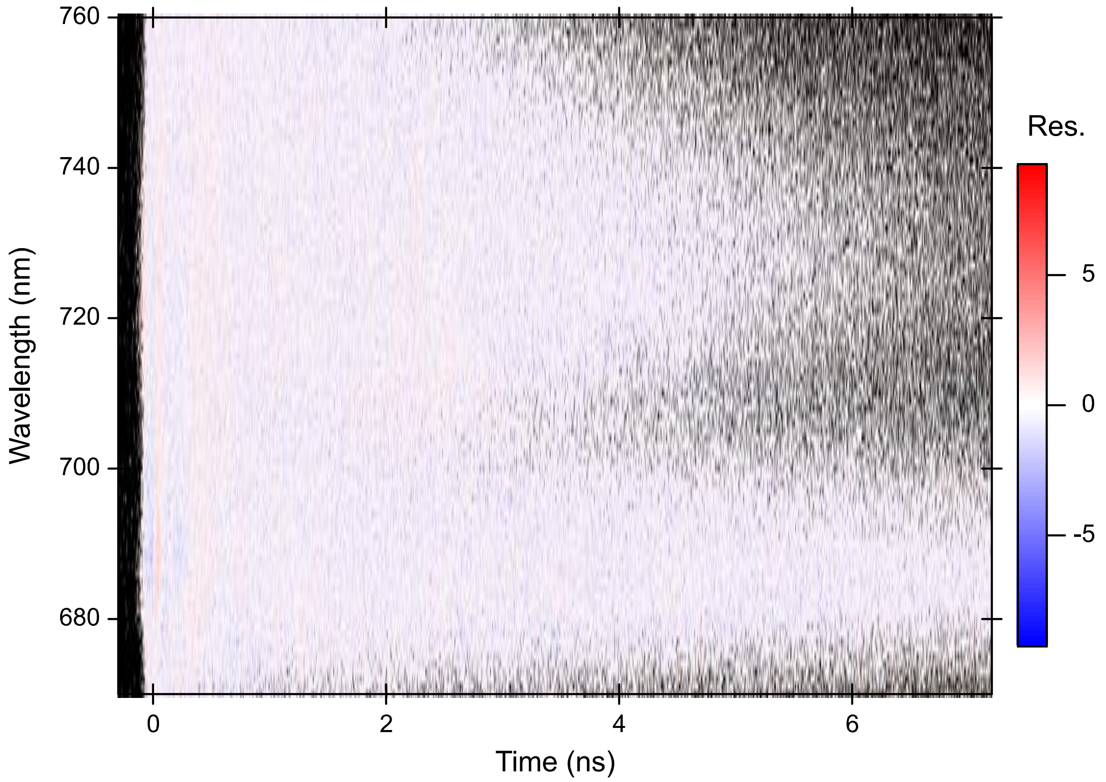


**Supplementary Fig. 11. Residual map of the global analysis.**

Residual (Res.) was defined as

$$Res. = \frac{I_m(t,\lambda) - I_c(t,\lambda)}{\sqrt{I_m(t,\lambda)}}.$$

Here, $I_m(t,\lambda)$ and $I_c(t,\lambda)$ are the measured and calculated data, respectively. Black dots indicate the points at which $I_m(t,\lambda)$ is 0.

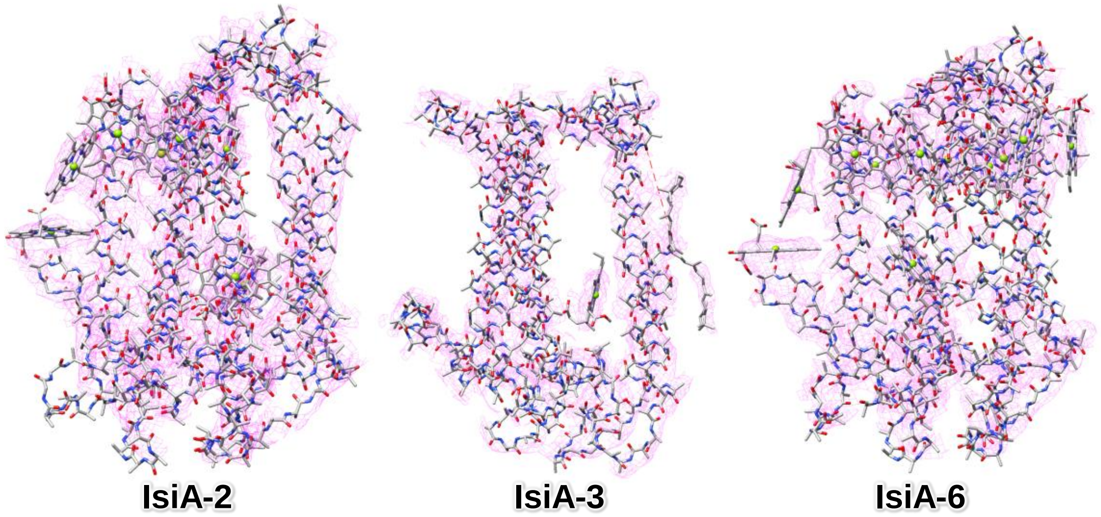


**Supplementary Fig. 12. Evaluation of the maps and model for IsiA-2, IsiA-3, and IsiA-6**
The densities for IsiA-2, IsiA-3, and IsiA-6 and their corresponding models are shown as magenta meshes and gray sticks, respectively, using the denoised map (see Methods).

**Supplementary Table 1. Cryo-EM data collection and structural analysis statistics.**

| | |
|---|---|
| Complex | PSI-IsiA |
| PDB ID | 7Y3F |
| EMDB ID | EMD-33593 |
| Data collection and processing | |
| Magnification | 100,000 |
| Voltage (kV) | 300 |
| Electron exposure ($e^{-}$/Å) | 40.8 |
| Defocus range (μm) | −1.8 to −0.8 |
| Pixel size (Å) | 0.495 |
| Symmetry imposed | C1 |
| Final particle images (no.) | 47,602 |
| Map resolution (Å) | 2.62 |
| FSC threshold | 0.143 |
| Refinement | |
| Initial model used (PDB code) | Homology modeling |
| Model resolution (Å) | 2.57 |
| FSC threshold | 0.5 |
| Map sharpening B factor (Å$^2$) | −45.4 |
| Model composition | |
| Non-hydrogen atoms | 39,463 |
| Protein | 24,774 |
| Ligand | 14,663 |
| Water | 26 |
| B factors (Å$^2$) | |
| Protein | 97.7 |
| Ligand | 145.3 |
| Water | 44.3 |
| R.m.s deviations | |
| Bond lengths (Å) | 0.028 |
| Bond angles (°) | 2.86 |
| Validation | |
| MolProbity score | 1.78 |
| Clashscore | 7.96 |
| Poor rotamers (%) | 0.36 |
| EMRinger score | 4.53 |
| Ramachandran plot | |
| Favored (%) | 95.02 |
| Allowed (%) | 4.21 |
| Disallowed (%) | 0.77 |

**Supplementary Table 2. Averaged *Q*-score in each subunit.**

| Subunit | Averaged *Q*-score | |
|---|---|---|
| | Postprocessed map | Denoised map |
| **PsaA** | 0.80 | 0.78 |
| **PsaB** | 0.79 | 0.78 |
| **PsaC** | 0.81 | 0.77 |
| **PsaD** | 0.76 | 0.74 |
| **PsaE** | 0.73 | 0.72 |
| **PsaF** | 0.77 | 0.75 |
| **PsaI** | 0.78 | 0.76 |
| **PsaJ** | 0.78 | 0.76 |
| **Unknown** | 0.74 | 0.73 |
| **PsaM** | 0.76 | 0.74 |
| **PsaX** | 0.73 | 0.72 |
| **IsiA2-1** | 0.57 | 0.60 |
| **IsiA-2** | 0.24 | 0.34 |
| **IsiA-3** | 0.22 | 0.33 |
| **IsiA1-4** | 0.30 | 0.40 |
| **IsiA1-5** | 0.34 | 0.43 |
| **IsiA-6** | 0.25 | 0.40 |

**Supplementary Table 3. Cofactors in the PSI-IsiA supercomplex.**

| Protein | Chlorophyll | Carotenoid | Lipid | Others |
|---|---|---|---|---|
| **PsaA** | 45 Chl *a*<br>1 Chl *a*' | 7 BCR | 2 LHG | 1 [4Fe-4S] cluster,<br>1 phylloquinone |
| **PsaB** | 41 Chl *a* | 7 BCR | 1 LMG<br>2 LHG | 1 phylloquinone |
| **PsaC** | - | - | - | 2 [4Fe-4S] cluster |
| **PsaD** | - | - | - | - |
| **PsaE** | - | - | - | - |
| **PsaF** | 1 Chl *a* | 1 BCR | - | - |
| **PsaI** | - | 2 BCR | - | - |
| **PsaJ** | 2 Chl *a* | 3 BCR | - | - |
| **Unknown** | 1 Chl *a* | - | - | - |
| **PsaM** | - | 1 BCR | - | - |
| **PsaX** | 1 Chl *a* | - | - | - |
| **IsiA2-1** | 17 Chl *a* | 5 BCR | - | - |
| **IsiA-2** | 8 Chl *a* | - | - | - |
| **IsiA-3** | 1 Chl *a* | 1 BCR | - | - |
| **IsiA1-4** | 10 Chl *a* | 1 BCR | - | - |
| **IsiA1-5** | 17 Chl *a* | 1 BCR | - | - |
| **IsiA-6** | 11 Chl *a* | - | - | - |
| **Total** | 156 | 29 | 5 | 5 |

BCR, $\beta$-carotene; Chl *a*, chlorophyll *a*; LMG, distearoylmonogalactosyl diglyceride; LHG, dipalmitoylphosphatidyl glycerol.

**Supplementary Table 4. IsiA proteins identified in the *Anabaena* PSI-IsiA structure and their RMSD values with IsiA1-5.**

| Protein | Gene | RMSD (Å)/Aligned Cα atoms |
|---|---|---|
| **IsiA2-1** | *isiA2* | 0.95/302 |
| **IsiA-2** | unidentified | 0.90/302 |
| **IsiA-3** | unidentified | 0.94/277 |
| **IsiA1-4** | *isiA1* | 0.56/326 |
| **IsiA1-5** | *isiA1* | 0.00/332 |
| **IsiA-6** | unidentified | 0.73/303 |

**Supplementary Table 5. Primers used in this study.**

| Primer | Sequence (5′- -3′) |
|---|---|
| RTisiA1-F | CCCAAGCTGCTCTGACAAC |
| RTisiA1-R | AAAATAAGTCCTTGCTCACCCA |
| RTisiA2-F | GAGAGTGCGTGTCCTAGAGG |
| RTisiA2-R | CCCGGACGATAAATTGGCAG |
| RTisiA3-F | AACTAGCCCGTTTCCAGACA |
| RTisiA3-R | TAACCTGACCACCACTACCC |
| RTisiA5-F | GGGCAGGAACAACTACCATC |
| RTisiA5-R | ACTTGACCACCAACACCTACT |
| RTrrn16S-F2 | GCAAGTCGAACGGTCTCTTC |
| RTrrn16S-R2 | GGTATTAGCCACCGTTTCCA |
